# Beneficial rhizobacteria and virus infection modulate the soybean metabolome and influence the feeding preferences of the virus vector *Epilachna varivestis*

**DOI:** 10.1101/2024.09.11.612434

**Authors:** Hannier Pulido, Kerry E. Mauck, Consuelo M. De Moraes, Mark C. Mescher

## Abstract

There is growing evidence that microbial plant symbionts shape interactions between plants and other organisms by modulating gene expression and metabolism. However, the detailed mechanisms mediating such effects are not well understood, particularly in systems where plants interact simultaneously with multiple symbionts and antagonists. In this study, we employed a multi-factorial design to explore the individual and combined effects of two plant-beneficial rhizobacteria (*Delftia acidovorans* and *Bradyrhizobium japonicum*) and a pathogen (*Bean pod mottle virus*: BPMV) on gene expression and metabolite production by soybean plants, as well as downstream effects on plant interactions with a beetle vector of BPMV *Epilachna varivestis*. Our results document microbial effects on basic metabolism and defense pathways, resulting in increased levels of primary metabolites and depletion of secondary metabolites. These changes are consistent with the observed feeding preferences of beetles for rhizobia-inoculated and virus-infected plants. Together, our results indicate that BPMV infection and rhizobacteria colonization cause dramatic changes in plant metabolites related to nutrition and defense, with significant consequences for an agriculturally important pathosystem.

## Introduction

Plant-associated microbes can alter biochemical pathways within their hosts in ways that affect ecological interactions among plants and other organisms. Beneficial rhizobacteria, for example, have been shown to modulate the production of primary and secondary plant metabolites, as well as the emission volatile organic compounds, with implications for the interactions of plants with herbivores, herbivore natural enemies, and vector-borne plant pathogens (Pineda et al. 2013; Pulido et al. 2019). Meanwhile, there is growing evidence that pathogens themselves can similarly alter plant phenotypes, frequently in ways that enhance transmission by arthropod vectors (Chesnais et al. 2020). Given that mutualistic and pathogenic symbionts can influence similar molecular and biochemical pathways (e.g., those influencing plant defense and nutritional quality for herbivores) but have potentially divergent fitness interests with respect to the alteration of plant phenotypes, questions arise regarding outcomes in the, likely typical, case where plants simultaneously engage with beneficial and pathogenic symbionts. However, few studies to date have explored such complex interactions. In this context, we investigate how these symbiotic interactions collectively shape the metabolic pathways of soybean plants and influence the foraging patterns of *Epilachna varivestis*, an important vector of BPMV. We aim to provide novel insights into the broader consequences of co-occurring symbiotic associations for plant ecology and agricultural pathosystems.

There is a substantial body of research documenting how viruses can alter host phenotypes in ways that influence insect vector behavior, typically enhancing transmission efficiency (Fereres and Moreno 2009; Ingwell et al. 2012; Mauck et al. 2018; Ray and Casteel 2022). For instance, infection of cucumber plants by *Chlorotic yellows virus* reduced defensive compounds, while increasing primary metabolites, such as amino acids and, which led to enhanced feeding (and presumably virus acquisition) by the whitefly vector *Bemisia tabaci* (Zhang et al. 2022). Similarly, Luan *et al*. (2013) found that infection by *Tomato yellow leaf curl china virus* triggered metabolite changes in tobacco plants, improving *B. tabaci* performance on this typically unfavorable host. Mauck *et al*. (2010) showed that *Cucumber mosaic virus* infection increased aphid attraction to infected squash plants but promoted rapid dispersal of the vectors—conditions conducive to the transmission of this non-persistently transmitted virus. Additionally, some studies have identified specific viral genes responsible for transmission-enhancing metabolic changes in host plants (Ray and Casteel 2022).

While viruses may modify host phenotypes in ways that promote their own transmission, beneficial soil-dwelling rhizobacteria frequently alter host-plant physiology in ways that enhance both bacterial and plant fitness (Berendsen et al. 2012; Zamioudis and Pieterse 2012). These modifications extend beyond root tissue and can have a systematic influence on the entire plant. This systemic effect can lead to changes in the levels of primary and secondary metabolites in ways that influence multi-trophic interactions among plants and other organisms (Pii et al. 2015; Grunseich et al. 2019). For example, Pulido *et al*. (2019) reported that co-colonization of soybean roots by a nitrogen-fixing bacterium and a plant-growth promoting rhizobacteria (PGPR) enhanced recruitment of parasitoid wasps to plants damaged by a specialist beetle herbivore. The current work builds upon that system, further exploring the interactions between rhizobacteria, soybean plants, and insect herbivores, demonstrating additional facets of this multi-trophic relationship.

Insight into the mechanisms underlying rhizobia effects on plant phenotypes has been provided by large-scale gene expression profiling approaches. These studies have revealed the molecular interplay between *Bradyrhizobium japonicum* and soybean roots as the bacteria establish and proliferate within root nodules (Libault et al. 2009). For instance, *B. japonicum* colonization alters the expression of genes involved in canonical and antioxidant-based plant defenses, as well as those involved in primary metabolism (Brechenmacher et al. 2008; Carvalho et al. 2013). Timecourse gene expression profiles further indicate that these effects are temporally dynamic and depend on the stage of the nodulation process (Libault et al. 2010). From a metabolomics perspective, *B. japonicum* nodulation has been shown to trigger rapid up-regulation of metabolites such as flavonoids, amino acids, and carbohydrates in soybean root hairs (Brechenmacher et al. 2010).

These studies demonstrate the power of transcriptomics and metabolomics in providing mechanistic insights into microbe-induced effects on plant phenotypes. When these approaches are employed in combination, they are particularly valuable for uncovering plant trait changes relevant to multi-trophic ecological interactions (Kang et al. 2018; Mhlongo et al. 2018; Castro-Moretti et al. 2020). However, this integrated approach has been underutilized in microbe-plant interaction studies, especially in exploring cocolonization by multiple symbionts, despite the likelihood that such interactions are widespread in nature.

In the present study, we address this research gap by examining co-colonization dynamics involving two beneficial microbes *B. japonicum* (**Bj**) and *Delftia acidovorans* (**Da**), alongside *Bean pod mottle virus* (**BPMV**) in soybean plants (*Glycine max* cv. Williams 82). We used enrichment pathway analysis, combining metabolomics and transcriptomics to identify signature genes and metabolites affected by these interactions (Fig. 1). To link these molecular insights to ecological outcomes, we conducted behavioral assays to assess the feeding preferences and larval performance of *Epilachna varivestis*, a beetle vector, in response to co-colonized plants. We hypothesize that the presence of beneficial rhizobacteria and BPMV induces distinct metabolic shifts in soybean, which in turn influence beetle feeding behavior, potentially affecting insect herbivory and overall plant performance.

**Figure 1:**
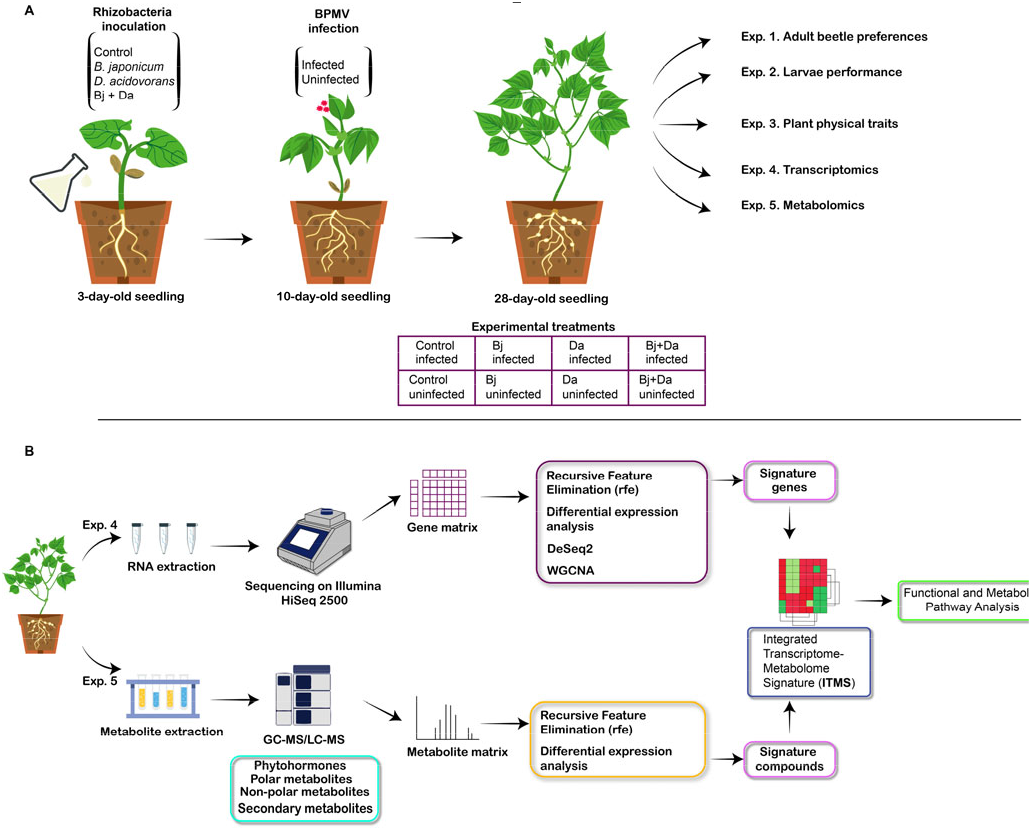
Experimental Design Overview. **A)** Rhizobacteria inoculation and virus infection protocols were used to generate experimental treatment plants. This design was applied across various experiments detailed in the Methods section. **B)** Integration of transcriptomics and metabolomics data. The Integrated Transcriptome-Metabolome Signature (ITMS) was created by merging identified features. Pathway analysis and group comparisons were then performed to elucidate the impacted metabolic processes.

## Results

### Effects on beetle performance

#### Both rhizobacteria inoculation and BPMV infection affect foraging and feeding preferences of adult beetles

In our dual-choice feeding experiment, we found distinct patterns of foraging activity. Uninfected control plants were more preferred by adult beetles than plants inoculated with either rhizobacteria species (binomial test, Bj proportion = 0.327, 95% CI [0.319, 0.335], p < 0.001; Da proportion = 0.996, 95% CI [0.986, 1.0], p < 0.001; Bj+Da proportion = 0.387, 95% CI [0.380, 0.394], p < 0.001; Fig. 2 top). However, in BPMV-infected plants, beetles preferred Bj- and Da-inoculated plants over control plants, while they foraged less on Bj+Da double-inoculated plants (binomial test, Bj proportion = 0.942, 95% CI [0.938, 0.945], p < 0.001; Da proportion = 0.484, 95% CI [0.479, 0.489], p = 0.001; Bj+Da proportion = 0.44, 95% CI [0.436, 0.444], p < 0.001; Fig. 2 middle). These results reveal an interaction between rhizobacteria inoculation and virus infection.

**Figure 2:**
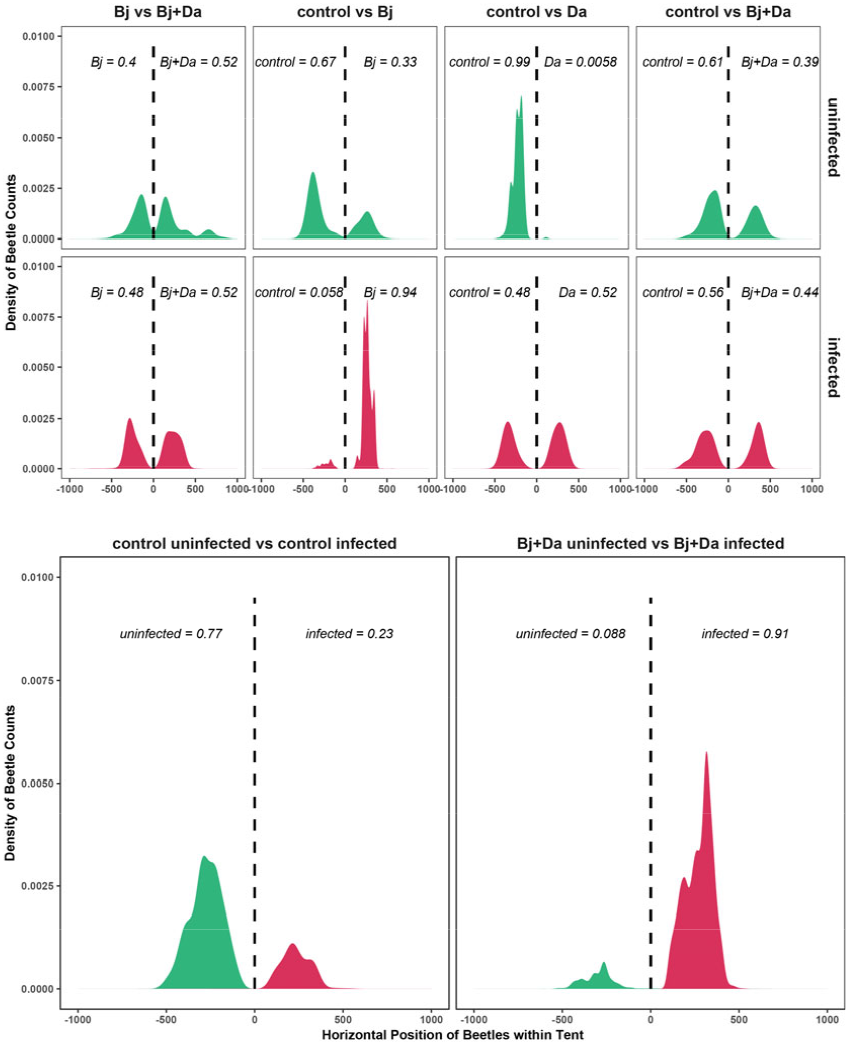
Adult Foraging Behavior in a Dual-Choice Test. Foraging behavior of beetles when rhizobacteria-inoculated plants were either uninfected (**top**) or infected with BPMV (**middle**). The **bottom** panel compares beetle foraging behavior between infected and uninfected plants for both Bj+Da inoculated plants and control plants. The x-axis represents the horizontal position of the beetles within the tent, and the density plots represent the time beetles spent at each plant. The vertical dashed lines indicate the midpoint of the tent. Each choice test was repeated twelve times with five different beetles per trial. Group probabilities are shown within the plots, and the presence of beetles on each side of the tent was compared using an exact binomial test. Uninfected treatments are represented in green, and infected treatments are shown in red. “Bj” = *Bradyrhizobium japonicum*, “Da” = *Delftia acidovorans*.

Further analysis showed that beetles preferred feeding on BPMV-infected Bj+Da-inoculated plants over uninfected Bj+Da plants (binomial test, proportion = 0.912, 95% CI [0.912, 0.913], p < 0.001). In contrast, without rhizobacteria, beetles fed more on uninfected plants compared to BPMV-infected ones (binomial test, proportion = 0.226, 95% CI [0.217, 0.236], p < 0.001; Fig. 2 bottom).

We also found that beetles fed more on Bj-inoculated plants compared to controls, regardless of BPMV infection (Fig. 3 top; one-way ANOVA, control vs Bj uninfected: F_1,22_= 7.153, p-value = 0.01; control vs Bj infected: F_1,12_= 6.419, p-value = 0.03; control vs Bj+Da uninfected: F_1,22_ = 7.223, p-value = 0.01; control vs Bj+Da infected: F_1,22_ = 20.272, p-value = 1.7×10^−4^). When offered a choice between uninfected and BPMV-infected plants with identical rhizobacteria treatments, beetles consistently preferred BPMV-infected plants (Fig. 3 bottom; one-way ANOVA, Bj+Da uninfected vs infected: F_1,38_ = 5.166, p-value = 0.028; control uninfected vs infected: F_1,22_ = 13.602, p-value = 0.001).

**Figure 3:**
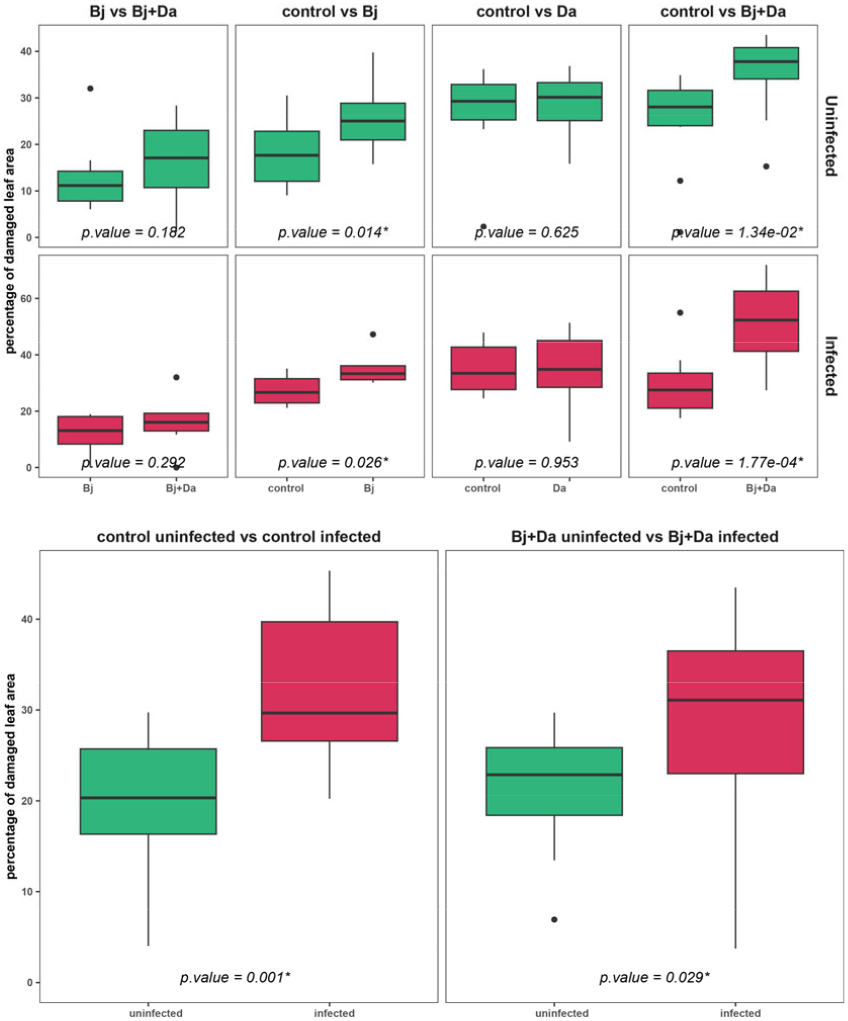
Adult Feeding Behavior in a Dual-Choice Test. Feeding behavior of beetles when rhizobacteria-inoculated plants were either uninfected (**top**) or infected with BPMV (**middle**). The **bottom** panels compare beetle feeding behavior between infected and uninfected plants for both Bj+Da inoculated plants and control plants. The y-axis represents the percentage of damaged leaf area. Uninfected treatments are represented in green, while infected treatments are shown in red. “Bj” = *Bradyrhizobium japonicum*, “Da” = *Delftia acidovorans*.

#### Both rhizobacteria inoculation and BPMV infection increase larvae gained weight

In our no-choice assay, beetle larvae gained significantly more weight on BPMV-infected plants (two-way ANOVA, F_1,46_ = 13.487, p-value = 0.001; Fig. 4). Rhizobacteria inoculation also had a significant effect (F_3,46_ = 5.355, p-value = 0.003; Fig. 4), though there was no significant interaction between BPMV infection and rhizobacteria (F_3,46_ = 0.12, p-value = 0.947). A post hoc Tukey test showed that larvae feeding on Bj+Da plants gained more weight than those feeding on control (p-value = 0.01; Fig. 4) or Da-inoculated plants (p-value = 0.02; Fig. 4).

**Figure 4:**
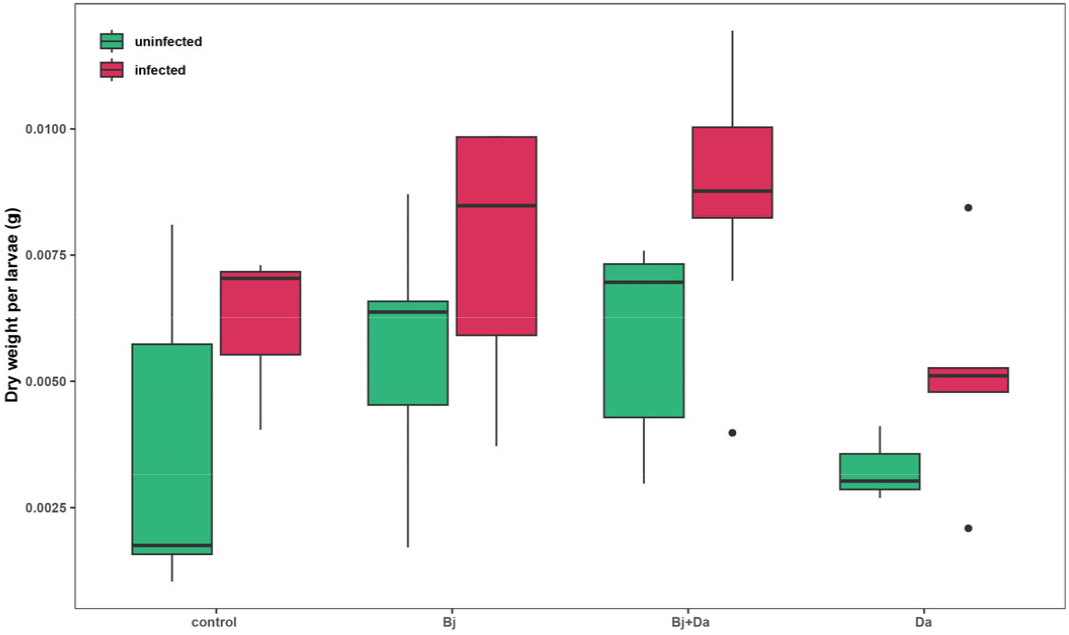
Larvae weight gain. Boxplots show the dry weight (g) gained per larva in a non-choice assay. Recently hatched larvae were placed on each plant and weighed when they reached the prepupal stage. Ten larvae were placed on each plant (n = 7). Dry weight was analyzed using a two-way ANOVA with rhizobacteria and virus infection as main factors. Uninfected treatments are represented in green, while infected treatments are shown in red. “Bj” = *Bradyrhizobium japonicum*, “Da” = *Delftia acidovorans*.

### Effects on plant physical traits

#### BPMV infection influences nodulation and shoot biomass but not leaf toughness

We did not observe significant effects of rhizobacteria inoculation or BPMV infection on leaf toughness (Fig. 5A, two-way ANOVA: rhizobacteria F_3,40_ = 1.28, p-value = 0.293, virus: F_1,40_ = 1.48, p-value = 0.230, interaction: F_3,40_ = 0.14 p-value = 0.933). BPMV-infected plants had significantly less biomass than uninfected plants, but there were no significant differences in plant shoot biomass between the rhizobacteria treatments and control plants (Fig. 5B, two-way ANOVA: rhizobacteria F_3,254_ = 1.789, p-value = 0.149; virus: F_1,254_ = 166.286, p-value < 0.001). Nodulation in Bj and Bj+Da plants was also significantly lower in BPMV-infected plants compared to control plants (Fig. 5C, two-way ANOVA: F_1,105_ = 28.785, p-value = 4.86×10^−7^). Using a simple linear regression model, we attempted to predict plant shoot biomass based on nodulation for Bj-uninfected, Bj-infected, Bj+Da-uninfected, and Bj+Da-infected independently. Two of the regression models showed a significant positive relationship between nodule weight and plant biomass (Fig. 5D): Bj-BPMV (F1,20 = 8.55, p = 0.008, R^2^ = 0.30) and Bj+Da-uninfected (F1,21 = 9.39, p = 0.005, R^2^ = 0.30). In contrast, the other two models did not reveal a significant relationship: Bj-uninfected (F1,29 = 0.96, p = 0.33, R^2^ = 0.03) and Bj+Da-BPMV (F1,6 = 0.69, p = 0.43, R^2^ = 0.10).

**Figure 5:**
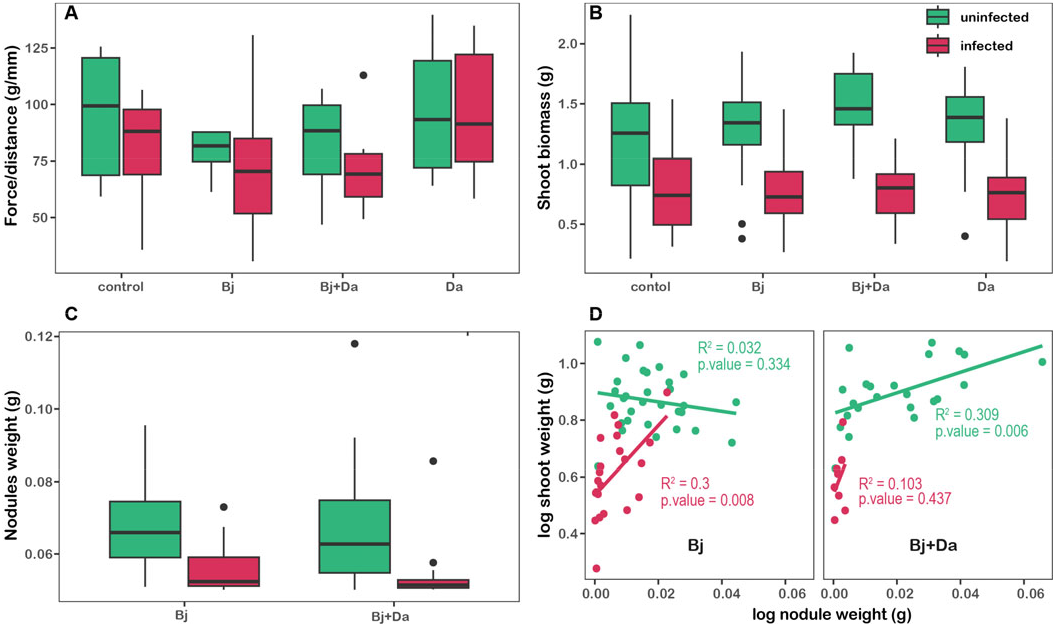
Plant physical traits. **A)** Leaf toughness, measured as the force required to penetrate the leaf tissue (force/distance in g/mm), was analyzed using a two-way ANOVA with rhizobacteria and BPMV infection as main factors. **B)** Shoot biomass for different rhizobacteria and virus treatments. **C)** Effect of BPMV infection on nodulation and D. acidovorans on nodulation. **D)** Regression analysis between nodule weight and shoot biomass for Bj and Bj+Da treatments. Uninfected treatments are represented in green, while infected treatments are shown in red. “Bj” = *Bradyrhizobium japonicum*, “Da” = *Delftia acidovorans*.

### Effects on plant metabolomics

#### Both Rhizobacteria inoculation and BPMV infection increase primary metabolite levels and reduce secondary metabolites

DAPC analysis revealed clear separation between uninfected and BPMV-infected samples, with Bj inoculation also strongly influencing group separation (Fig. 6A). Linear models identified 88 metabolites with significant changes due to rhizobacteria inoculation (allBacteria = combined Bj, Bj+Da, and Da treatments) in uninfected plants (65 accumulated and 23 depleted) (Fig. 7A). This number increased to 99 metabolites in plants that were both rhizobacteria-inoculated and BPMV-infected (75 accumulated and 24 depleted) (Fig. 7B). Specifically, in rhizobacteria-inoculated BPMV-infected plants, 10 metabolites (mainly carbohydrates and amino acids) were significantly accumulated, while 19 metabolites (mostly flavonoids) were significantly depleted (Fig. 7C).

**Figure 6:**
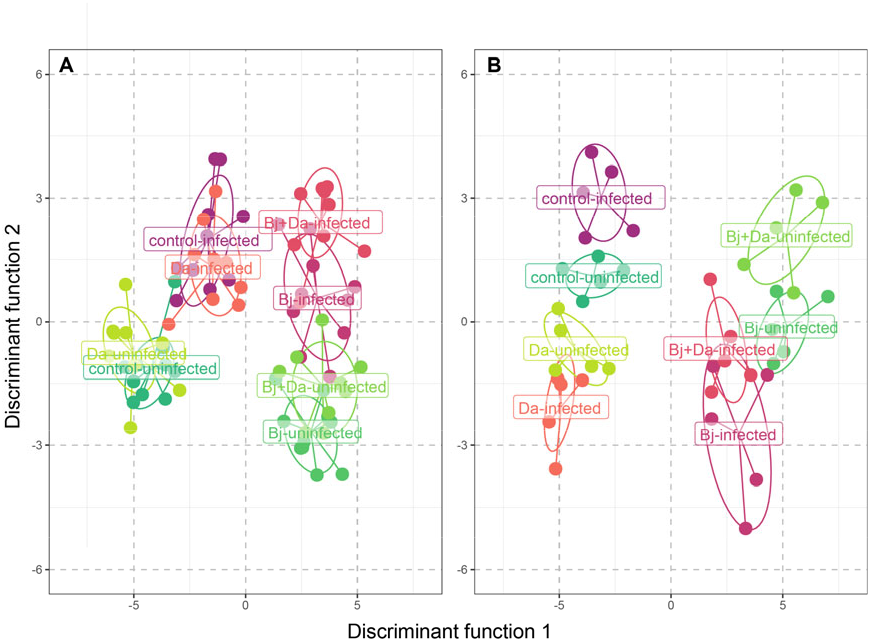
Group Separation Using Discriminant Analysis of Principal Components (DAPC). **A)** Clustering of samples based on metabolite abundance data, plotted along Discriminant Function 1 and Discriminant Function 2. Each point represents a sample, color-coded by treatment. Ellipses indicate clusters, highlighting the separation between treatments. **B)** Clustering of samples based on RNA-Seq normalized counts data. Uninfected treatments are represented in shades of green, while infected treatments are shown in shades of red. “Bj” = *Bradyrhizobium japonicum*, “Da” = *Delftia acidovorans*.

**Figure 7:**
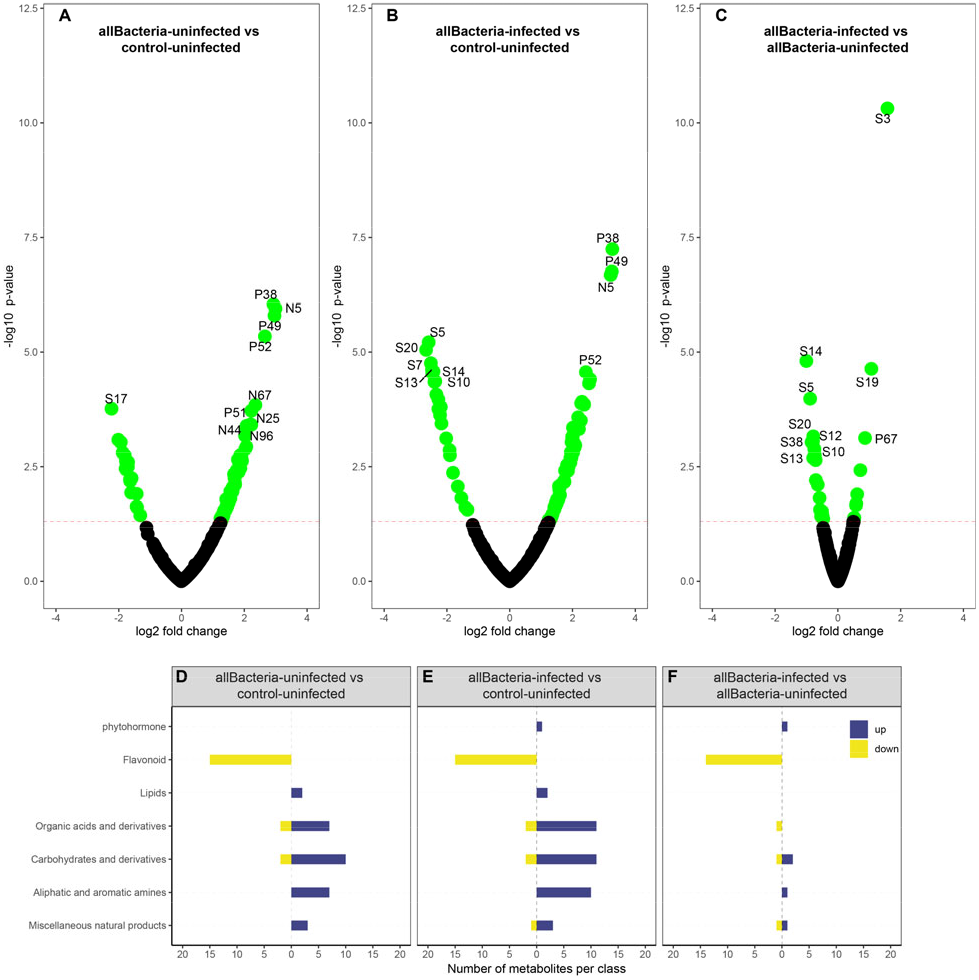
Modulation of metabolite profiles by rhizobacteria colonization and BPMV infection in soybean plants. The **top** panel shows differentially accumulated metabolites across three contrasts: **A)** All bacteria uninfected treatments compared to control uninfected. **B)** All bacteria, BPMV-infected treatments compared to control uninfected. **C)** All bacteria, BPMV-infected treatments compared to all bacteria uninfected treatments. Significant metabolites (p-value < 0.05) are highlighted as green dots, while non-significant metabolites (p-value > 0.05) are shown as black dots. The **lower** panel displays the number of differentially accumulated metabolites (DAMs) for the three contrasts, categorized as follows: 1) Organic acids and derivatives: carboxylic acid, fatty acid, organic acid, amide, peptide. 2) Carbohydrates and derivatives: carbohydrate, monosaccharide, sugar alcohol, glycoside. 3) Lipids: fat, fatty alcohol, lipid. 4) Aliphatic and aromatic amines: amine, amino acid, indole. 5) Miscellaneous natural products: acyloin, alcohol, aldehyde, glucosinolate, inorganic acid, pyranone, terpenoid, tetracycline. (Compound IDs are listed in supplementary table S2.3.). **all Bacteria** = *B. japonicum, B. japonicum + D. acidovorans*, and *D. acidovorans*.

Comparisons between individual rhizobacteria treatments in uninfected versus BPMV-infected plants (Figs. S3.2-S3.3, File S5.1) and between mixed uninfected and BPMV-infected treatments (Fig. S3.4, File S5.1) revealed 71 metabolites significantly changed in Bj+Da-inoculated uninfected plants (48 accumulated and 23 depleted) and 69 in Bj+Da-infected plants (27 accumulated and 42 depleted). BPMV infection altered 13 metabolites in Bj+Da plants (9 accumulated and 4 depleted) and 22 metabolites in control plants (21 accumulated and 1 depleted). Notably, both BPMV infection and rhizobacteria inoculation reduced flavonoid production and increased organic acids, carbohydrates, and amino acids.

Furthermore, rhizobacteria inoculation of uninfected plants led to a more than 5-fold increase in several metabolites, including beta-alanine, glyceric acid, and pyruvic acid, compared to control plants. The presence of both rhizobacteria and BPMV further increased certain metabolites while reducing flavonoid levels (Table 1, Fig. 7C, S3.4, File S5.1).

**Table 1.**
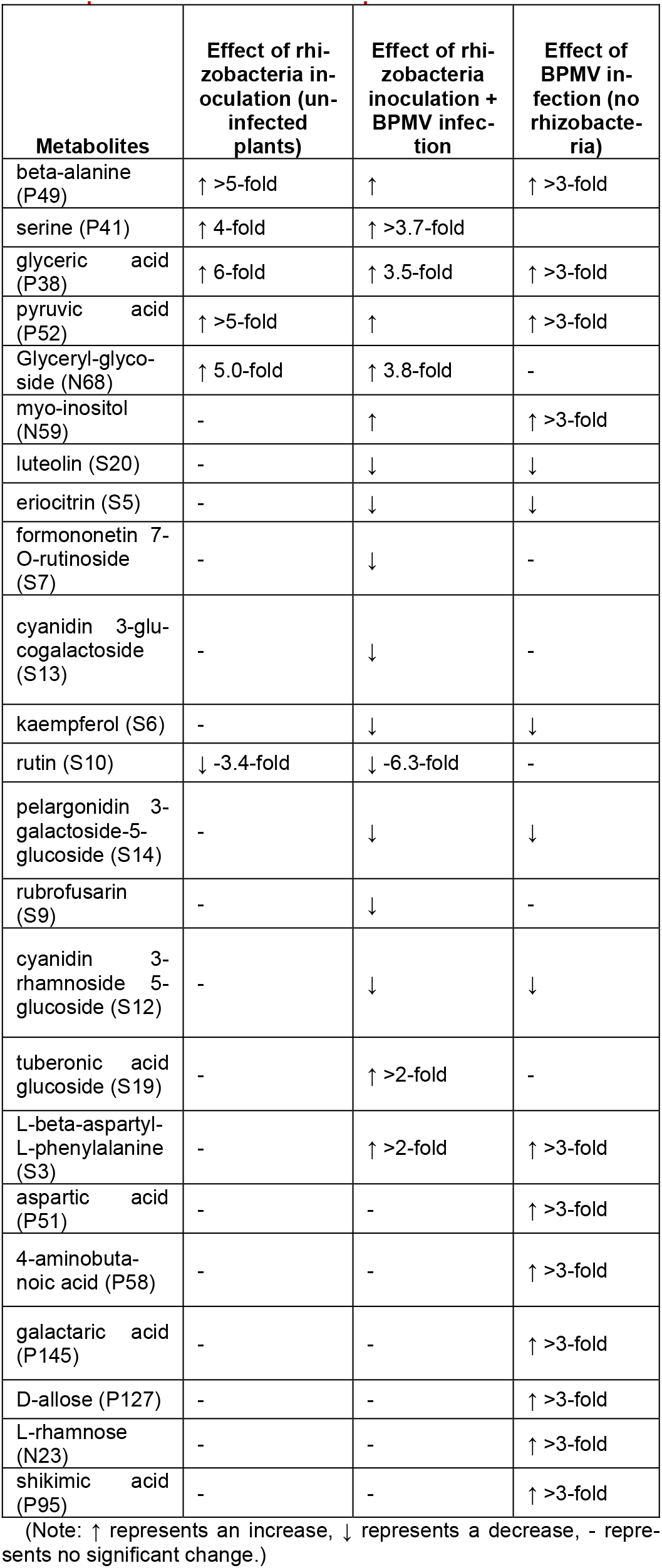
Metabolite changes upon rhizobacteria inoculation and BPMV infection. The compound code used in the graphs is provided in parentheses after each compound name.

| Metabolites | Effect of rhizobacteria inoculation (uninfected plants) | Effect of rhizobacteria inoculation + BPMV infection | Effect of BPMV infection (no rhizobacteria) |
| --- | --- | --- | --- |
| beta-alanine (P49) | ↑ >5-fold | ↑ | ↑ >3-fold |
| serine (P41) | ↑ 4-fold | ↑ >3.7-fold |  |
| glyceric acid (P38) | ↑ 6-fold | ↑ 3.5-fold | ↑ >3-fold |
| pyruvic acid (P52) | ↑ >5-fold | ↑ | ↑ >3-fold |
| Glyceryl-glycoside (N68) | ↑ 5.0-fold | ↑ 3.8-fold | - |
| myo-inositol (N59) | - | ↑ | ↑ >3-fold |
| luteolin (S20) | - | ↓ | ↓ |
| eriocitrin (S5) | - | ↓ | ↓ |
| formononetin 7-O-rutinoside (S7) | - | ↓ | - |
| cyanidin 3-glucogalactoside (S13) | - | ↓ | - |
| kaempferol (S6) | - | ↓ | ↓ |
| rutin (S10) | ↓ -3.4-fold | ↓ -6.3-fold | - |
| pelargonidin 3-galactoside-5-glucoside (S14) | - | ↓ | ↓ |
| rubrofusarin (S9) | - | ↓ | - |
| cyanidin 3-rhamnoside 5-glucoside (S12) | - | ↓ | ↓ |
| tuberonic acid glucoside (S19) | - | ↑ >2-fold | - |
| L-beta-aspartyl-L-phenylalanine (S3) | - | ↑ >2-fold | ↑ >3-fold |
| aspartic acid (P51) | - | - | ↑ >3-fold |
| 4-aminobutanoic acid (P58) | - | - | ↑ >3-fold |
| galactaric acid (P145) | - | - | ↑ >3-fold |
| D-allose (P127) | - | - | ↑ >3-fold |
| L-rhamnose (N23) | - | - | ↑ >3-fold |
| shikimic acid (P95) | - | - | ↑ >3-fold |

(Note: ↑ represents an increase, ↓ represents a decrease, - represents no significant change.)

#### BPMV infection decreases the levels of some rhizobacteria-induced metabolites

BPMV infection of rhizobacteria-inoculated plants led to a reduced concentration of some metabolites that were previously accumulated. Notably, metabolites such as glyceric acid, glyceryl-glycoside, and serine showed decreased fold changes in BPMV-infected plants compared to their uninfected rhizobacteria-inoculated counterparts. Additionally, the depletion of flavonoid compounds was more pronounced following BPMV infection (Table 1).

Metabolite compounds were grouped based on their chemical properties. Our analysis confirmed that both BPMV infection and rhizobacteria inoculation caused significant decreases in flavonoid compounds, while increasing levels of organic acids, carbohydrates, and amino acids compared to control uninfected plants (Fig. 7D, E). Interestingly, BPMV infection of rhizobacteria-inoculated plants also led to a reduction in the abundance of primary metabolites, including lipids, organic acids, carbohydrates, and amino acids (Fig. 7F).

### Effects on plant transcriptomics

#### BPMV and rhizobacteria treatments shift soybean gene expression patterns

We analyzed changes in soybean gene expression profiles induced by BPMV infection and rhizobacteria inoculation using RNA-Seq followed by DAPC (see Methods). As shown in Fig. 6B, both rhizobacteria inoculation (Bj, Da, and Bj+Da) and BPMV infection strongly affected soybean gene expression.

To gain detailed insights, we identified **signature genes**—sets of genes whose expression changed coordinately under different conditions—based on differential expression (DESeq), importance (RFE), and gene-module membership (WGCNA). We performed multiple pairwise comparisons, assessing individual and combined effects of rhizobacteria and virus infection (Fig. S3.5, Table S2.2, and Files S5.1-S5.4). We found a high number of signature genes upon inoculation with Bj and Bj+Da, and fewer with Da, indicating that the nitrogen-fixing symbiont Bj has a stronger effect on soybean gene expression compared to the growthpromoting Da. Additionally, fewer rhizobacteria-induced signature genes were observed in control plants than in BPMV-infected plants (Fig. S3.6 and File S5.2). For instance, 90 upregulated and 21 downregulated genes were identified in uninfected Bj+Da versus Bj plants, compared to 860 upregulated and 1530 downregulated genes in BPMV-infected plants (Fig. S3.5, File S5.2), highlighting the strong impact of BPMV on soybean gene regulation and microbial responses.

Fig. 8 displays the top 10 signature genes induced by rhizobacteria inoculation (All Bacteria), BPMV infection, or their combination. Among these, SODB2 (iron-superoxide dismutase), ELF4A, and ELF4B (EARLY FLOWERING 4a and 4b) were upregulated in both uninfected and BPMV-infected plants (Figs. 8A, 7B). BPMV infection of rhizobacteria-inoculated plants resulted in upregulation of all top 10 signature genes: LOC100791680 (NAC domain-containing protein 73), LOC100788906 (AAA-ATPase At3g50940),

**Figure 8:**
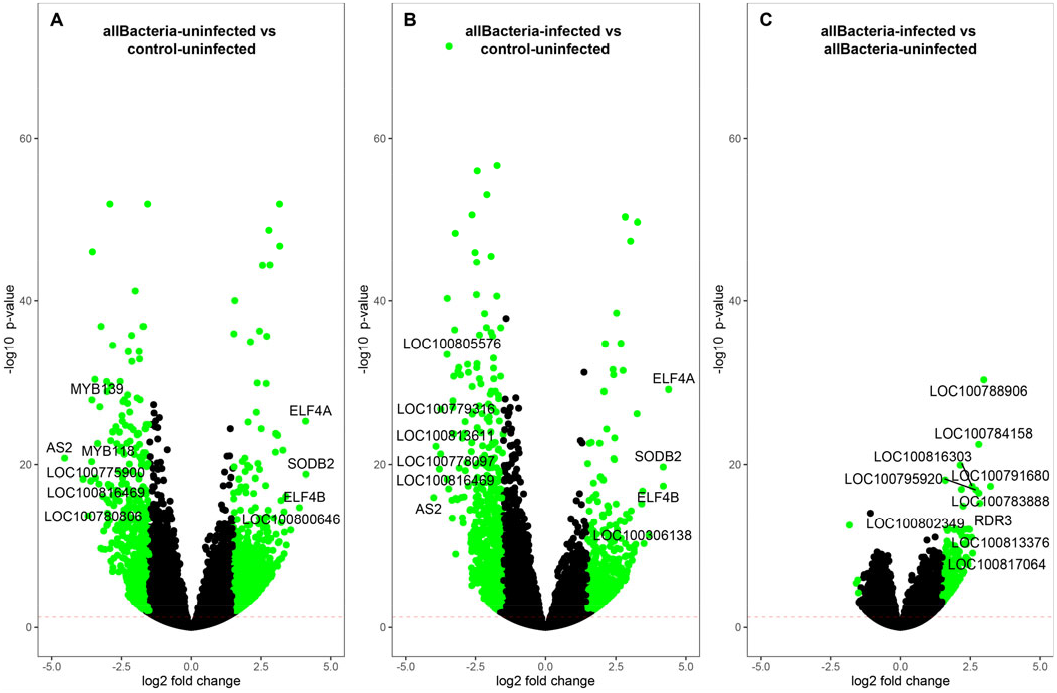
Volcano plots depicting gene expression profiles in soybean plants under rhizobacteria colonization and BPMV infection. Signature genes identified through DESeq analysis, recursive feature elimination, and WGCNA are labeled. Significant genes (log2 fold change > 1.5 and adjusted p-value < 0.01) are highlighted in green, while non-significant genes are shown in black. The y-axis represents -log10 p-values, and the x-axis shows log2 fold changes for each gene. **A)** Gene expression in all uninfected rhizobacteria treatments compared to control uninfected. **B)** Gene expression in all BPMV-infected rhizobacteria treatments compared to control uninfected. **C)** Gene expression in all BPMV-infected rhizobacteria treatments compared to all uninfected rhizobacteria treatments. **all Bacteria** = *B. japonicum, B. japonicum + D. acidovorans*, and *D. acidovorans*.

RDR3 (RNA-dependent RNA polymerase), LOC100783888 (NAC domain-containing protein 73), LOC100784158 (probable WRKY transcription factor 23-like), LOC100795920 (NAC domain-containing protein 73), LOC100817064 (pumilio homolog 2), LOC100816303 (TMV resistance protein N-like), LOC100813376 (NAC domain-containing protein 87), and LOC100802349 (ureide permease 1) (Fig. 8C). Additional comparisons are detailed in Figs. S3.6-S3.8 and File S5.2.

#### Integration of signature genes and metabolites reveals changes in defense, energy, and nutrition pathways

Using the full transcriptomics dataset, we identified 48 co-expression modules (MEs) containing between 127 and 9,735 genes, totaling 41,540 genes (Fig. 9A). Two modules, ME6 and ME1, were significantly regulated in response to rhizobacteria inoculation regardless of BPMV infection, with ME6 genes strongly upregulated and ME1 genes downregulated (Figs. 9B-C and File S5.4). Conversely, two other modules, ME2 and ME15, were significantly downregulated in rhizobacteria-inoculated plants either in the absence of BPMV (ME2) or in its presence (ME15) (Figs. 9B-C and File S5.4). Additionally, ME21 was significantly upregulated by BPMV infection, independent of rhizobacteria inoculation (Figs. 9C-D, 8E, and File S5.4), while modules ME20 and ME7 were regulated by BPMV infection only in the presence of rhizobacteria, with ME7 genes upregulated and ME20 genes downregulated (Figs. 9B-C and File S5.4). The top six genes for each of these modules are listed in Table S2.4, and the expression profiles of key modules ME6, ME1, and ME7 are shown in Figs. 9E-G.

**Figure 9:**
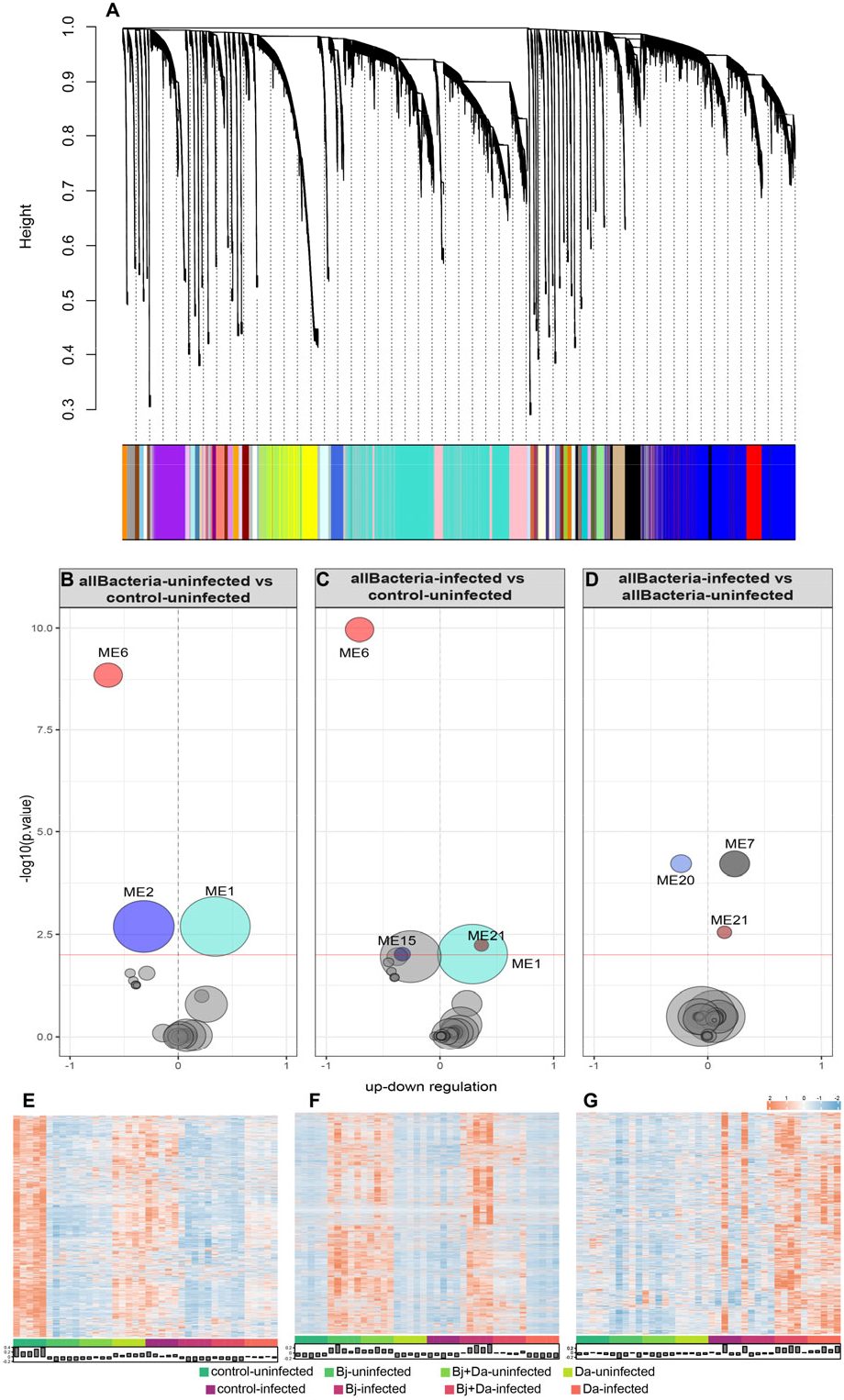
Weighted Gene Co-expression Network Analysis (WGCNA) analysis of whole leaf transcriptomics on soybean plants after rhizobacteria inoculation and BPMV-infection. **A)** Transcripts were grouped into 49 modules based on co-expression patterns. The dendrogram shows the hierarchical relationships and dissimilarities between modules, with branch height indicating the degree of dissimilarity. Each module is assigned a unique color by WGCNA, displayed in the color strip at the bottom. Module sizes vary from 127 to 9,735 genes, with size proportional to the number of genes. **Middle panel**: Significant modules detected by WGCNA across three contrasts: **B)** All uninfected rhizobacteria treatments vs. control uninfected. **C)** All BPMV-infected rhizobacteria treatments vs. control uninfected. **D)** All BPMV-infected rhizobacteria treatments vs. all uninfected rhizobacteria treatments. Significant modules (adjusted p-value < 0.01) are shown in their respective WGCNA colors, while non-significant modules are depicted in grey. Module size is proportional to the number of genes assigned. **Bottom panel**: Gene expression profiles for the most significant modules: **E)** Module 6, **F)** Module 1, **G)** Module 7. Uninfected treatments are shaded in green, and BPMV-infected samples in red. Bar plots represent module eigengenes, summarizing gene expression patterns for each sample in each module. **all Bacteria** = *B. japonicum, B. japonicum + D. acidovorans*, and *D. acidovorans*.

To gain further insights, we performed a gene/metabolite enrichment analysis (GAGE) to identify significantly perturbed KEGG pathways in response to these treatments. Rhizobacteria inoculation in uninfected plants upregulated pathways related to carbohydrate, lipid, nucleotide, and amino acid metabolism, while downregulating carotenoid biosynthesis (gmx00906), MAPK signaling (gmx04016), and flavonoid biosynthesis (gmx00941) (Figs. 10A, D, G; Fig. S3.9A; File S5.5).

**Figure 10:**
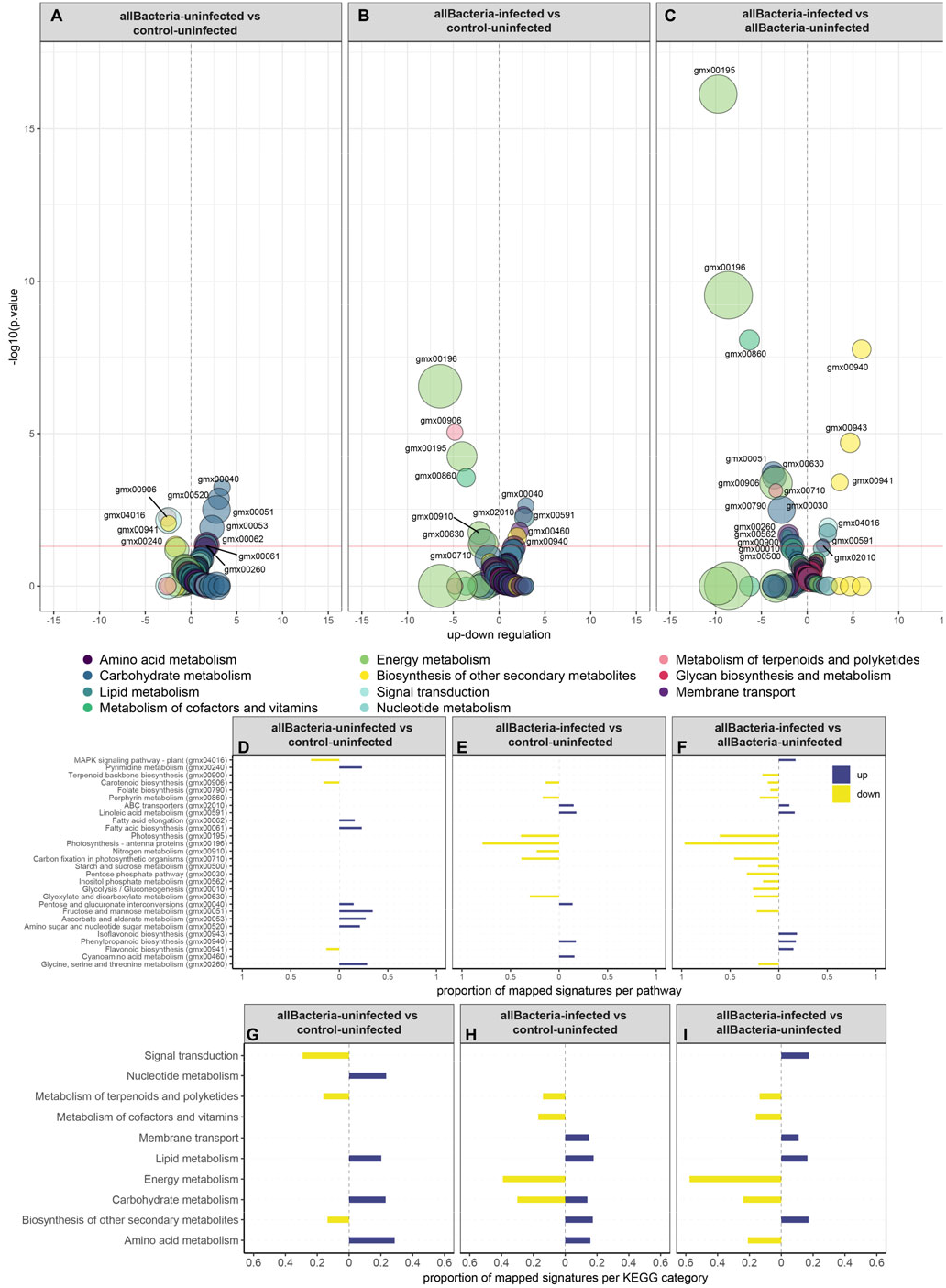
Gene-set Enrichment (GAGE) and KEGG pathway analysis of the Integrated Transcriptome-Metabolome Signature (ITMS). The **top panel** shows the distribution of KEGG pathways across three contrasts: **A)** All uninfected rhizobacteria treatments vs. control uninfected. **B)** All BPMV-infected rhizobacteria treatments vs. control uninfected. **C)** All BPMV-infected rhizobacteria treatments vs. all uninfected rhizobacteria treatments. Significantly enriched pathways identified through GAGE (p-value < 0.05) are labeled by their KEGG pathway ID. Colored bubbles indicate the KEGG category for each pathway, and bubble size represents the proportion of signature genes and metabolites mapped to that pathway. Pathways are color-coded based on their KEGG category. The **middle panel** shows the proportion of signature genes and metabolites mapped to significant KEGG pathways for each contrast. The **bottom panel** illustrates the proportion of mapped signatures within KEGG categories for the same contrasts. **all Bacteria** = *B. japonicum, B. japonicum + D. acidovorans*, and *D. acidovorans*.

Interestingly, in rhizobacteria-inoculated, BPMV-infected plants, there was a reduction in the upregulation of carbohydrate metabolism pathways and a strong downregulation of pathways related to energy metabolism, nitrogen metabolism, cofactors, vitamins, and terpenoids and polyketides (Figs. 10B, E, H; File S5.5).

Further analysis revealed that BPMV infection significantly perturbed pathways associated with secondary metabolite biosynthesis, signal transduction, membrane transport, and energy, carbohydrate, and amino acid metabolism in rhizobacteria-inoculated plants (Bj, Bj+Da, and Da). Notably, phenylpropanoid biosynthesis (gmx00940), isoflavonoid biosynthesis (gmx00943), flavonoid biosynthesis (gmx00941), and MAPK signaling (gmx04016) were the most significantly upregulated pathways, indicating their enrichment due to BPMV infection (Fig. S3.9C). In contrast, photosynthesis (gmx00195), photosynthesis antenna proteins (gmx00196), porphyrin metabolism (gmx00860), fructose and mannose metabolism (gmx00051), and glyoxylate and dicarboxylate metabolism (gmx00630) were the most significantly downregulated pathways, highlighting a strong impact on energy and carbohydrate metabolism (Figs. 10C, F, I; Fig. S3.9C; and File S5.5).

## Discussion

Beneficial rhizobacteria and viral pathogens are known to play pivotal roles in shaping plant phenotypes and influencing trophic interactions with insects. Using a combined metabolite profiling and transcriptomics approach, we investigated soybean’s response to both BPMV infection and rhizobacteria inoculation, with implications for the behavior of *Epilachna varivestis*, a semi-specialist beetle herbivore and BPMV vector. Our results highlight the profound impact of simultaneous colonization by *Bradyrhizobium japonicum* and *Delftia acidovorans*, which disrupts core metabolic and defense pathways. This disruption led to an increase in primary metabolites such as amino acids and fatty acids, alongside a depletion of flavonoids in both rhizobacteria-colonized uninfected and BPMV-infected plants. Notably, similar metabolic shifts were observed in plants infected with BPMV alone. These changes likely contribute to the altered beetle feeding preferences observed when compared to rhizobacteria-free, uninfected plants. In summary, our findings address key questions about how co-colonization by beneficial rhizobacteria and BPMV infection alters soybean metabolism and subsequently impacts the feeding behavior of *E. varivestis*, which could play a role in virus transmission.

### Effects on beetle performance

Larvae that fed on BPMV-infected plants displayed significantly greater weights than those feeding on healthy plants (Fig. 4), consistent with previous research on *E. varivestis* feeding behavior on BPMV-infected *Phaseolus vulgaris* (Musser et al. 2003). This relationship may enhance the beetle’s fitness, as faster growth and maturation could benefit the virus by promoting its transmission to additional plant hosts. A similar trend of heightened growth was observed in larvae fed on *B. japonicum*-inoculated plants. Although rhizobacteria do not directly benefit from beetle feeding, this outcome can be attributed to the enhanced nutritional value and reduced defensive compounds present in rhizobacteria-inoculated plants. This aligns with previous findings in other beneficial interactions between rhizobacteria and herbivores (Pineda et al. 2012; Dean et al. 2014).

Adult beetles also exhibited a strong preference for feeding on both rhizobacteria-inoculated plants and BPMV-infected plants (Fig. 3). In the context of a vector herbivore, virus-induced changes in plant metabolites can drive vector feeding behavior and facilitate virus dispersal to new hosts (Mauck et al. 2010; Luan et al. 2013). This mechanism encourages vector dispersal after feeding and virus acquisition, as vectors that remain on infected plants are less likely to spread the virus to other hosts (Mauck et al. 2010).

We found evidence supporting this dispersal behavior in our system in the absence of rhizobacteria. While adult beetles fed more on BPMV-infected plants (leading to virus acquisition), they foraged more on uninfected plants (Fig. 2, bottom panel). This behavior could enhance virus dispersal to healthy plants. However, when plants were inoculated with both rhizobacteria species, an opposite foraging preference emerged (Fig. 2, bottom panel). Adult beetles continued to feed more on rhizobacteria-inoculated, BPMV-infected plants, but they also spent significantly more time foraging on these plants, showing less inclination to disperse. This suggests that adult beetles may be less likely to disperse from BPMV-infected plants if these plants are also simultaneously inoculated by *B. japonicum* and *D. acidovorans*.

This finding aligns with a previous study (Pulido et al. 2019) using the same experimental setup, where BPMV infection was shown to reduce the release of volatile compounds in soybean plants. Volatile emissions are crucial for attracting herbivores, and the reduction in volatiles due to BPMV infection may diminish the plants’ attractiveness to beetles. Consequently, herbivores already present on infected plants may be more inclined to move to uninfected plants after acquiring the virus. Notably, rhizobacteria inoculation appears to influence this pattern, as evidenced by the increased attraction of beetles to ***Bj+Da***-infected plants. This could be due to rhizobacteria restoring volatile emissions, as suggested by Pulido *et al*. (2019).

### Effects on mechanisms of interactions

The metabolic changes in soybean plants after BPMV infection, as well as the suppression of defense pathways after rhizobacteria inoculation and nodule formation, suggest a complex interplay between plant defense and symbiosis mechanisms. Although we cannot fully pinpoint specific rhizobacteria and virus-induced changes responsible for *Epilachna varivestis* feeding and foraging preferences, our combined transcriptomic and metabolomic approaches do allow us to generate some hypotheses like the increased feeding being explained by reduced defenses and increased primary metabolites and the possibility that the activation of growth-related pathways in response to rhizobacteria colonization might come at the cost of defense mechanisms.

### BPMV infection induces defense and flavonoid biosynthesis genes yet lowers flavonoid levels

Our metabolic profiles indicate that BPMV is the primary driver of beetle feeding preferences and foraging behavior. BPMV infection significantly depleted flavonoid compounds, including eriocitrin, luteolin, and kaempferol, in soybean plants (Fig. 7 and Table S2.3), despite the upregulation of flavonoid biosynthesis and other defense-related genes. These changes in flavonoid biosynthesis have not been previously reported in soybeans following BPMV infection, though the induction of defense pathways is well documented in response to pests and necrotrophic fungi (Dastmalchi et al. 2017). Flavonoids act as repellents or toxins to insects (Treutter 2006), and isoflavonoid phytoalexins are known to deter *E. varivestis* feeding (Hart et al. 1983). Therefore, the depletion of these compounds in BPMV-infected plants likely increases beetle feeding preference, encouraging vector herbivores to feed on infected plants, aiding virus transmission.

Despite the reduction in flavonoid levels, BPMV infection triggered the upregulation of pathways associated with phenylpropanoid, isoflavonoid, and flavonoid biosynthesis, along with other defense-related secondary metabolites. Similar upregulation of these pathways in response to viral infection-induced stress has been documented in previous studies (Jiang et al. 2021). The apparent contradiction— lower flavonoid levels despite increased expression of biosynthesis genes—could be due to the plant’s defense response being counteracted by viral effectors through posttranscriptional mechanisms.

The upregulation of phenylpropanoid biosynthesis and pentose and glucuronate interconversion pathways in BPMV-infected plants suggests that the phenylpropanoid pathway may compensate for flavonoid depletion. This pathway is crucial for lignin synthesis, providing structural support and defense against pathogens and herbivores. Elevated levels of L-beta-aspartyl-L-phenylalanine and phenylalanine ammonia-lyase (PAL) in BPMV-infected plants (Files S5.1 and S5.2) further support this hypothesis, suggesting that phenylalanine is being diverted into the phenylpropanoid pathway rather than the flavonoid pathway. The downregulation of purine metabolism in BPMV-infected plants aligns with studies showing that viral infections can disrupt nucleotide metabolism in plants (Yue et al. 2018).

Additionally, BPMV infection led to notable changes in soybean metabolite concentrations. Infected plants showed increased levels of sugars and organic acids, such as D-glucuronic acid, D-fructose, and pyruvic acid. This is consistent with previous research reporting higher amino acid and carbohydrate levels in BPMV-exposed soybean plants (Peñaflor et al. 2016), though it contrasts with findings of reduced amino acid concentrations in a more recent study (Smith et al. 2017). Furthermore, BPMV infection increased concentrations of amino acids such as glycine, aspartic acid, and 4-aminobutanoic acid, all of which play roles in plant defense. Their accumulation may indicate an activated defense response in BPMV-infected plants (Rojas et al. 2014; Mur et al. 2017).

### *Bradyrhizobium japonicum* colonization redirects soybean plant resources from defense mechanisms towards nodule development and symbiotic nitrogen fixation

In soybean plants inoculated with rhizobacteria, the depletion of flavonoid compounds, the downregulation of carotenoid biosynthesis and MAPK signaling pathways, along with reduced levels of phenylalanine ammonia-lyase (PAL) transcripts, suggest a shift away from defense mechanisms. These pathways are crucial for producing defense-related compounds and signaling molecules. For instance, carotenoids play a role in protecting plants from herbivores by acting as deterrents or toxins, and their suppression could make soybean plants more susceptible to insect attack. Recent studies support this, showing that carotenoid depletion can weaken plant defenses, leading to increased herbivory (Jones et al. 2019; Chen et al. 2021). This correlates with our findings, where *E. varivestis* fed more on plants colonized by *B. japonicum* (Fig. 3A), causing increased damage. These results emphasize the complexity of plant-microbe interactions and the trade-offs between growth and defense.

Conversely, in plants co-inoculated with both *B. japonicum* and *D. acidovorans* (Bj+Da), we observed an increase in primary metabolites like amino acids, carboxylic acids, and fatty acids, alongside upregulated pathways related to amino sugar and nucleotide sugar metabolism, and pentose and glucuronate interconversions. This suggests a resource shift towards growth and development, particularly nodule formation and symbiotic nitrogen fixation, driven by *B. japonicum* inoculation. These results align with previous studies, which report activation of growth-related pathways and concurrent downregulation of defense pathways in plants inoculated with nodule-forming rhizobacteria (Brechenmacher et al. 2008). While recent research has challenged the idea that growth-related pathways always detract from defense (Afzal et al. 2019), our findings indicate that in the case of *B. japonicum* colonization, significant resource reallocation occurs, resulting in enhanced plant growth at the cost of decreased defense and increased herbivory

## Conclusion

Our study demonstrates that soybean leaf metabolite production is intricately regulated by the combined effects of beneficial rhizobacteria and BPMV infection, with significant implications for the feeding behavior of *Epilachna varivestis*. The results highlight a complex interaction between plant defense mechanisms and symbiosis, emphasizing the importance of considering co-occurring microbes when evaluating virus-induced host phenotypes in natural environments.

These findings carry broader implications for understanding plant-microbe interactions and their role in agricultural ecosystems. While beneficial microorganisms are increasingly being used in sustainable agriculture to enhance plant growth and defense against pests and diseases, our research suggests that their influence on plant metabolism can be multifaceted and may lead to unintended consequences for herbivore behavior and fitness. Thus, a more nuanced understanding of these interactions is crucial for developing sustainable agricultural practices that effectively balance plant growth, defense, and ecosystem health.

Future research into the role of co-occurring microbes in virus-induced plant phenotypes will help us unravel the complexities of plant defense and symbiosis. This knowledge is key in informing strategies to enhance plant health and productivity, especially in the context of environmental challenges faced by modern agriculture.

## Materials and Methods

### Rhizobacteria and BPMV culture conditions

We used a multi-factorial design to investigate the effects of single, dual, and triple colonization by the plant-growth-promoting rhizobacteria *Delftia acidovorans* (Da), the nitrogen-fixing root symbiont *Bradyrhizobium japonicum* (Bj), and the virus pathogen *Bean pod mottle virus* (BPMV) on soybean plants (Fig. 1, Table S2.1). This design allowed us to test various combinations of these treatments. The number of replicates varied depending on the specific experiments described below. Rhizobacteria cultures were isolated from a commercial product (BrettYoung) and kept for long-term storage under sterile conditions at -80°C as 30% glycerol stocks. A BPMV strain that was collected in Ohio (US) was used to mechanically inoculate soybean plants in V1 (see Supplementary Methods S1.1 for details).

### Plant growth and experimental design

Soybean seeds were sterilized for 5 minutes in a 10% sodium hypochlorite solution and washed with copious amounts of ultrapure water. Seeds were germinated in a sterilized growing medium (Premier Pro-mix without mycorrhiza, Griffin Supplies) autoclaved at 120°C for 40 min. Three-day old seedlings were transplanted to individual 500 ml pots containing the same growing medium and inoculated with rhizobacteria. One week after rhizobacteria inoculation, seedlings were infected with BPMV or mock-inoculated according to the multi-factorial design described in Fig.1 and Table S2.1. Starting from the V1 stage, plants received 50 ml of a modified Hoagland’s nutrient solution three times per week. Plants inoculated with Bj alone or in combination with Da received the same nutrient solution but without nitrogen, as nodulation is inhibited by the presence of nitrates in the soil (Carroll and Mathews 2018). This strategy for supplementing nutrient solution to non-rhizobium inoculated plants has also been employed in our previous work (Pulido et al. 2019) and by other studies exploring similar systems (Silva et al. 2013; Dean et al. 2014). All plants were kept in an insect-free growth chamber at 25°C (day), 23°C (night), a L16:D8 photoperiod and at 70% of relative humidity (see Supplementary Methods S1.2 for details).

### Beetle colonies

Colonies of the Mexican bean beetle (*Epilachna varivestis*, Coccinellidae), a significant herbivore of legumes and a known pest of soybeans, were initially provided by the New Jersey Department of Agriculture’s Phillip Alampi Beneficial Insect Laboratory. *E. varivestis* were maintained on *Phaseolus vulgaris* plants in an incubator at 25°C and a L16:D8 photoperiod. One day prior to the bioassays, beetles were moved to the greenhouse where behavioral experiments took place to allow them to habituate to the new conditions.

### Adult beetle preferences and larval performance

We assessed larval performance using no-choice assays focusing on rhizobacteria and virus treatments. Ten newly hatched larvae were placed on each plant (n = 7) and allowed to feed until reaching the prepupal stage. After approximately two weeks, prepupa were removed, dried at 50°C for two days and total dry weight was recorded and divided by the number of remaining larvae to calculate the average weight. Plants were covered with a transparent mesh net to restrict larvae from moving to other plants. We analyzed the average weight using ANOVA with rhizobacteria and virus infection as main factors.

We quantified adult feeding and foraging preferences using a dual-choice assay where two plants from different treatments were placed inside a fine-mesh cage (Fig. S3.1, supplementary video S.4). During the test, five beetle adults were released within the tent and allowed to freely forage for 24 hours. The combined activity (feeding or foraging behavior) of the five beetles within a tent was considered an individual replicate, as adult beetles do not tend to aggregate on individual host plants. Each choice test was repeated 12 times over the course of 8 days. After the test was completed, beetles were removed and the leaf damage was quantified using Photoshop CC software and analyzed in a two-way ANOVA with rhizobacteria and virus infection as main factors. The following dual-choice pairings were performed for uninfected and infected plants: **a**. Bj vs. Bj+Da, **b**. Bj vs control, **c**. Da vs Control, and **d**. Bj+Da vs Control. Additionally, we tested the following mixed BPMV treatments: **e**. Bj+Da-BPMV vs control-uninfected.

During the assay, we also recorded the beetle activity inside a tent using three DSLR cameras, the time-lapse videos were analyzed using Kinovea software and beetle movement was converted into a plain text file with three spatial coordinates. We restricted the analyses to the space coordinates where the plants were present. Each text file that represented the movement of five beetles inside a tent was considered a test replicate for a total of three replicates per paired-choice test during the time of the experiment. This spatial analysis gives insight about foraging preferences of adult beetles and propensity of movements relevant for BPMV transmission. Beetle’s foraging behavior was evaluated using an exact binomial test assuming a null distribution probability of 0.5 to forage on either side of the tent. In this test, beetles were treated as independent samples as adult beetles were observed to be non-gregarious.

### Plant physical traits

Before grinding the leaf tissue for metabolite extraction, samples were weighed, and total leaf biomass was recorded for each plant within a treatment. After harvesting the aboveground tissue, roots were thoroughly washed and nodules were collected and placed separately inside paper envelopes, dried at 50 °C for 48 h and the total dry biomass of the nodules was measured for each plant (nodules are only present in plants inoculated with *Bj*, but control and Da plants were checked for possible cross-contamination).

A TA-XT2i Texture Analyser (Texture Technologies Corp., Scarsdale, NY, USA) with a 3-mm diameter spherical probe was used to measure the force (N) it takes to penetrate soybean leaf tissue using six replicates per treatment (rhizobacteria treatments crossed with BPMV-infected plants, one leaf per plant was used). The test speed was 1 mm/s. Leaf toughness is a physical trait that has been implicated as an important factor in direct defenses against herbivores (Marquis et al. 2002; Malishev and Sanson 2015). In our model system we tested whether beneficial rhizobacteria and virus infection played a role in promoting differences in leaf toughness and whether this trait might also explain the differences observed in feeding behavior.

Leaf toughness (Force/distance(g/mm)), total shoot biomass, nodule biomass and leaf toughness were analyzed separately in a two-way ANOVA with rhizobacteria and virus infection as main factors. We used a regression analysis to assess the effect of nodulation on shoot biomass for each rhizobacteria and virus interaction.

### Phytohormone extraction and analysis

Plant tissue was harvested at 3-4 weeks of age (V4 stage), weighed (100-150 mg) and flash frozen in liquid nitrogen (n = 10 total replications per treatment). For phytohormone extraction, we used a protocol modified from Schmelz *et al*. (2004). Briefly, frozen tissue was pulverized in a mill grinder (Spex Certiprep GenoGrinder 2000) under liquid nitrogen. The homogenized material received a mixture of 100 ng of internal, isotopic standards for cis-OPDA, jasmonic acid (Tokyo Chemical Industry Co. (Tokyo, Japan)), salicylic acid (Campro Scientific (Berlin, Germany)), abscisic acid, linoleic acid, and linolenic acid. Phytohormone extraction took place in an acidic buffer solution of water, propanol and HCL. The organic phase was derivatized to methyl esters using Trimethylsilyldiazomethane (Sigma-Aldrich, St. Louis, MO). The remaining solvent was evaporated, and vials were heated to 200 C while headspace was pulled across an adsorbent trap for two minutes (30mg Super-Q). Super-Q traps were eluted with 150 μl dichloromethane and the eluate was analyzed by GC-MS in single ion mode with isobutene chemical ionization. Final concentrations of free phytohormones were quantified in the MassHunter software relative to recovery of the internal standards. Amounts of phytohormones were corrected by the original fresh weight of the sample. Data were integrated with other metabolite information for multivariate analysis.

### Metabolite extraction and analysis

Immediately after harvesting the tissue for phytohormones, remnant leaf tissue per plant was collected in paper bags and frozen in liquid nitrogen (n = 10 total replications per treatment). All tissue was lyophilized for 72h and ground to powder in a GenoGrinder. Once all the tissue per plant was homogenized, 10 +/- 0.6 mg of dried tissue was weighed into a 4.0 ml glass vial and processed with a multi-phasic extraction protocol for polar, non-polar and secondary metabolites (Broeckling et al. 2005) (see Methods S1.3 for details on sample and data processing).

We developed a Shiny app to provide an interactive and user-friendly interface to visualize the effect of rhizobacteria and virus infection on soybean metabolites. The app allows users to select the metabolite of interest and compare its abundance between the two treatments. It is available at https://hannier.shinyapps.io/Metabolites_shinyApp/

### RNA extraction

To assess the effect of rhizobacteria co-inoculation and BPMV on soybean gene expression, we harvested plant leaf tissue from a separate set of untouched undamaged plants at the V4 stage. Tissue sections with weights of 100-200 mg were obtained, and the tissue was flash frozen in liquid nitrogen. The homogenized material was extracted for total RNA using a mirVana miRNA isolation kit (Ambion, Austin, TX), with modifications suggested by Peña-Llopis and Brugarolas (2013). We combined equal amounts of three individual total RNA samples with similar quality and concentration to obtain a pooled replicate sample for each treatment, resulting in a total of five biological pooled replicates per treatment (n = 40). We used a Nanodrop Spectrophotometer (NanoDrop Technologies, Wilmington, DE) to determine RNA concentration and assessed RNA quality using the RNA Integrity Number (RIN) method, which is widely used for RNA quality control (Mueller et al. 2004), in an Agilent 2100 Bioanalyzer (Agilent Technologies, Santa Clara, CA).

### RNA-Seq library construction and sequencing

To prepare the barcoded libraries for the 40 RNA samples, 100 ng of total RNA was used with a TruSeq Stranded mRNA Library Kit, following the manufacturer’s instructions. The concentration of each library was determined by qPCR, and an equimolar pool of the 40 barcoded libraries was created. Sequencing was performed on an Illumina HiSeq 2500 in Rapid Run mode, using 150-nucleotide single-read sequencing. We conducted three consecutive sequencing runs to achieve the desired number of reads per sample. Raw sequence reads are available from the NCBI read archive under the accession number GSE244001

### Mapping and processing of RNA-seq reads

After base calling, adapter content and poor-quality reads were removed from the samples using *trimmomatic* (Bolger et al. 2014). The samples were then mapped to the reference genome of Glycine max Version 2.0 using *tophat2*, utilizing genome and annotation files obtained from GenBank (ftp://ftp.ncbi.nlm.nih.gov/genomes/Glycine_max/Assembled_chromosomes/seq/). Coverage files and a count matrix were generated with *bedtools*. The rlog-normalized feature counts matrix was then utilized for further statistical analysis, gene enrichment analysis, and pathway mapping.

### Transcriptomics and metabolomics data integration

To identify the metabolic processes affected by the interaction of beneficial rhizobacteria and BPMV infection on soybean, we integrated transcriptomic and metabolomic data using a functional and metabolic pathway analysis, as depicted in the flowchart (Fig. 1).

To simplify comparisons and account for similarities in gene expression and metabolite abundance, we grouped all uninfected rhizobacteria treatments (Bj, Bj+Da, and Da) into one category (“**all Bacteria-uninfected**”) and all BPMV-infected rhizobacteria treatments into another (“**all Bacteria-infected**”). This allowed us to compare these groups with the control-uninfected treatment and identify significant differences across analyses. See Methods S1.4 for details.

## Data availability

The complete results of the gene enrichment analysis (GAGE), linear models (limma), differential gene expression analysis (DESeq), and weighted correlation network analysis (WGCNA) are provided as Excel files in the supplementary information. Additional supplementary methods, tables, and figures are available in the ETHZ repository https://www.research-collection.ethz.ch/handle/20.500.11850/692860

The source code and data used for the analyses in this paper can be accessed in the following repository: https://gitlab.ethz.ch/hannierp/beetles_metabolomics.git

Raw sequence reads are available in the NCBI Sequence Read Archive (SRA) under the accession number GSE244001.

## Acknowledgements

The authors thank Heike Betz for her assistance with chemical analyses. Thomas Dorsey at the New Jersey Department of Agriculture’s Phillip Alampi Beneficial Insect Laboratory kindly provided us with beetles as needed, and Rene Mabon of the BrettYoung company for supplying the rhizobacterial inoculants. We are grateful to Istvan Albert and Aswathy Sebastian for their help with the initial processing and mapping of the transcriptomic data.

## Author contributions

H.P., K.E.M., M.C.M. and C.D.M. designed the study; H.P. carried out the experiments, collected and analyzed the data and drafted the initial manuscript. All authors contributed significantly to revisions and gave final approval for publication.

## References

Afzal I, Shinwari ZK, Sikandar S, Shahzad S. 2019. Plant beneficial endophytic bacteria: Mechanisms, diversity, host range and genetic determinants. Microbiol Res. 221:36–49. doi:10.1016/j.micres.2019.02.001.

Barros De Carvalho GA, Batista JSS, Marcelino-Guimarães FC, Costa Do Nascimento L, Hungria M. 2013. Transcriptional analysis of genes involved in nodulation in soybean roots inoculated with Bradyrhizobium japonicumstrain CPAC 15. BMC Genomics. 14(1):153. doi:10.1186/1471-2164-14-153.

Berendsen RL, Pieterse CMJ, Bakker PAHM. 2012. The rhizosphere microbiome and plant health. Trends Plant Sci. 17(8):478–486. doi:10.1016/j.tplants.2012.04.001.

Bolger AM, Lohse M, Usadel B. 2014. Trimmomatic: a flexible trimmer for Illumina sequence data. Bioinformatics. 30(15):2114–2120. doi:10.1093/bioinformatics/btu170.

Brechenmacher L, Kim M-Y, Benitez M, Li M, Joshi T, Calla B, Lee MP, Libault M, Vodkin LO, Xu D, et al. 2008. Transcription Profiling of Soybean Nodulation by Bradyrhizobium japonicum. Mol Plant-Microbe Interactions®. 21(5):631–645. doi:10.1094/MPMI-21-5-0631.

Brechenmacher L, Lei Z, Libault M, Findley S, Sugawara M, Sadowsky MJ, Sumner LW, Stacey G. 2010. Soybean Metabolites Regulated in Root Hairs in Response to the Symbiotic Bacterium Bradyrhizobium japonicum. Plant Physiol. 153(4):1808–1822. doi:10.1104/pp.110.157800.

Broeckling CD, Huhman DV, Farag MA, Smith JT, May GD, Mendes P, Dixon RA, Sumner LW. 2005. Metabolic profiling of Medicago truncatula cell cultures reveals the effects of biotic and abiotic elicitors on metabolism. J Exp Bot. 56(410):323–336. doi:10.1093/jxb/eri058.

Carroll BJ, Mathews A. 2018. Nitrate inhibition of nodulation in legumes, Molecular biology of symbiotic nitrogen fixation. CRC Press.

Castro-Moretti FR, Gentzel IN, Mackey D, Alonso AP. 2020. Metabolomics as an Emerging Tool for the Study of Plant–Pathogen Interactions. Metabolites. 10(2):52. doi:10.3390/metabo10020052.

Chen L, Sun D, Zhang X, Shao D, Lu Y, An Y. 2021. Transcriptome analysis of yellow passion fruit in response to cucumber mosaic virus infection. Sharma RK, editor. PLOS ONE. 16(2):e0247127. doi:10.1371/journal.pone.0247127.

Chesnais Q, Caballero Vidal G, Coquelle R, Yvon M, Mauck K, Brault V, Ameline A. 2020. Post-acquisition effects of viruses on vector behavior are important components of manipulation strategies. Oecologia. 194(3):429–440. doi:10.1007/s00442-020-04763-0.

Dastmalchi M, Chapman P, Yu J, Austin RS, Dhaubhadel S. 2017. Transcriptomic evidence for the control of soybean root isoflavonoid content by regulation of overlapping phenylpropanoid pathways. BMC Genomics. 18(1):70. doi:10.1186/s12864-016-3463-y.

Dean J, Mescher M, De Moraes C. 2014. Plant Dependence on Rhizobia for Nitrogen Influences Induced Plant Defenses and Herbivore Performance. Int J Mol Sci. 15(1):1466–1480. doi:10.3390/ijms15011466.

Fereres A, Moreno A. 2009. Behavioural aspects influencing plant virus transmission by homopteran insects. Virus Res. 141(2):158–168. doi:10.1016/j.virusres.2008.10.020.

Grunseich JM, Thompson MN, Aguirre NM, Helms AM. 2019. The Role of Plant-Associated Microbes in Mediating Host-Plant Selection by Insect Herbivores. Plants. 9(1):6. doi:10.3390/plants9010006.

Hart SV, Kogan M, Paxton JD. 1983. Effect of soybean phytoalexins on the herbivorous insects mexican bean beetle and soybean looper. J Chem Ecol. 9(6):657–672. doi:10.1007/BF00988774.

Ingwell LL, Eigenbrode SD, Bosque-Pérez NA. 2012. Plant viruses alter insect behavior to enhance their spread. Sci Rep. 2(1):578. doi:10.1038/srep00578.

Jiang L, Wu P, Yang L, Liu C, Guo P, Wang H, Wang S, Xu F, Zhuang Q, Tong X, et al. 2021. Transcriptomics and metabolomics reveal the induction of flavonoid biosynthesis pathway in the interaction of Stylosanthes-Colletotrichum gloeosporioides. Genomics. 113(4):2702–2716. doi:10.1016/j.ygeno.2021.06.004.

Jones P, Garcia BJ, Furches A, Tuskan GA, Jacobson D. 2019. Plant Host-Associated Mechanisms for Microbial Selection. Front Plant Sci. 10:862. doi:10.3389/fpls.2019.00862.

Kang W, Zhu X, Wang Y, Chen L, Duan Y. 2018. Transcriptomic and metabolomic analyses reveal that bacteria promote plant defense during infection of soybean cyst nematode in soybean. BMC Plant Biol. 18(1):86. doi:10.1186/s12870-018-1302-9.

Libault M, Joshi T, Takahashi K, Hurley-Sommer A, Puricelli K, Blake S, Finger RE, Taylor CG, Xu D, Nguyen HT, et al. 2009. Large-Scale Analysis of Putative Soybean Regulatory Gene Expression Identifies a Myb Gene Involved in Soybean Nodule Development. Plant Physiol. 151(3):1207–1220. doi:10.1104/pp.109.144030.

Luan J, Yao D, Zhang T, Walling LL, Yang M, Wang Y, Liu S. 2013. Suppression of terpenoid synthesis in plants by a virus promotes its mutualism with vectors. Irwin R, editor. Ecol Lett. 16(3):390–398. doi:10.1111/ele.12055.

Malishev M, Sanson GD. 2015. Leaf mechanics and herbivory defence: How tough tissue along the leaf body deters growing insect herbivores. Austral Ecol. 40(3):300–308. doi:10.1111/aec.12214.

Marquis RJ, Lill JT, Piccinni A. 2002. Effect of plant architecture on colonization and damage by leaftying caterpillars of Quercus alba. Oikos. 99(3):531–537. doi:10.1034/j.1600-0706.2002.11897.x.

Mauck KE, Chesnais Q, Shapiro LR. 2018. Evolutionary Determinants of Host and Vector Manipulation by Plant Viruses. In: Advances in Virus Research. Vol. 101. Elsevier. p. 189–250. [accessed 2024 Sep 11]. https://linkinghub.elsevier.com/retrieve/pii/S0065352718300101.

Mauck KE, De Moraes CM, Mescher MC. 2010. Deceptive chemical signals induced by a plant virus attract insect vectors to inferior hosts. Proc Natl Acad Sci. 107(8):3600–3605. doi:10.1073/pnas.0907191107.

Mhlongo MI, Piater LA, Madala NE, Labuschagne N, Dubery IA. 2018. The Chemistry of Plant–Microbe Interactions in the Rhizosphere and the Potential for Metabolomics to Reveal Signaling Related to Defense Priming and Induced Systemic Resistance. Front Plant Sci. 9:112. doi:10.3389/fpls.2018.00112.

Mueller O, Lightfoot S, Schroeder A. 2004. RNA integrity number (RIN)-standardization of RNA quality control. Agil Appl Note.:1–8.

Mur LA, Simpson C, Kumari A, Gupta AK, Gupta KJ. 2017. Moving nitrogen to the centre of plant defence against pathogens. Ann Bot. 119(5):703–709.

Musser RO, Hum-Musser SM, Felton GW, Gergerich RC. 2003. Increased larval growth and preference for virus-infected leaves by the Mexican bean beetle, Epilachna varivestis mulsant, a plant virus vector. J Insect Behav. 16:247–256. doi:10.1023/A:1023919902976.

Peñaflor MFG, Mauck KE, Alves KJ, Moraes CM, Mescher MC. 2016. Effects of single and mixed infections of bean pod mottle virus and soybean mosaic virus on host plant chemistry and host-vector interactions. Funct Ecol. 30:1648–1659. doi:10.1111/1365-2435.12649.

Peña-Llopis S, Brugarolas J. 2013. Simultaneous isolation of high-quality DNA, RNA, miRNA and proteins from tissues for genomic applications. Nat Protoc. 8:2240–2255. doi:10.1038/nprot.2013.141.

Pii Y, Mimmo T, Tomasi N, Terzano R, Cesco S, Crecchio C. 2015. Microbial interactions in the rhizosphere: beneficial influences of plant growth-promoting rhizobacteria on nutrient acquisition process. A review. Biol Fertil Soils. 51:403–415. doi:10.1007/s00374-015-0996-1.

Pineda A, Soler R, Weldegergis BT, Shimwela MM, Loon JJ, Dicke M. 2013. Non-pathogenic rhizobacteria interfere with the attraction of parasitoids to aphid-induced plant volatiles via jasmonic acid signalling. Plant Cell Environ. 36:393–404. doi:10.1111/j.1365-3040.2012.02581.x.

Pineda A, Zheng S -J., Van Loon JJA, Dicke M. 2012. Rhizobacteria modify plant–aphid interactions: a case of induced systemic susceptibility. Plant Biol. 14(1):83–90. doi:10.1111/j.1438-8677.2011.00549.x.

Pulido H, Mauck KE, De Moraes CM, Mescher MC. 2019. Combined effects of mutualistic rhizobacteria counteract virus-induced suppression of indirect plant defences in soya bean. Proc R Soc B Biol Sci. 286(1903):20190211. doi:10.1098/rspb.2019.0211.

Ray S, Casteel CL. 2022. Effector-mediated plant–virus–vector interactions. Plant Cell. 34(5):1514–1531. doi:10.1093/plcell/koac058.

Rojas CM, Senthil-Kumar M, Tzin V, Mysore KS. 2014. Regulation of primary plant metabolism during plant-pathogen interactions and its contribution to plant defense. Front Plant Sci. 5. doi:10.3389/fpls.2014.00017. [accessed 2024 Sep 11]. http://jour-nal.frontiersin.org/article/10.3389/fpls.2014.00017/abstract.

Schmelz EA, Engelberth J, Tumlinson JH, Block A, Alborn HT. 2004. The use of vapor phase extraction in metabolic profiling of phytohormones and other metabolites. Plant J. 39(5):790–808. doi:10.1111/j.1365-313X.2004.02168.x.

Silva LR, Pereira MJ, Azevedo J, Mulas R, Velazquez E, González-Andrés F, Valentão P, Andrade PB. 2013. Inoculation with Bradyrhizobium japonicum enhances the organic and fatty acids content of soybean (Glycine max (L.) Merrill) seeds. Food Chem. 141(4):3636–3648. doi:10.1016/j.foodchem.2013.06.045.

Smith CM, Gedling CR, Wiebe KF, Cassone BJ. 2017. A Sweet Story: Bean pod mottle virus Transmission Dynamics by Mexican Bean Beetles (Epilachna varivestis). Genome Biol Evol. 9(3):714–725. doi:10.1093/gbe/evx033.

Treutter D. 2006. Significance of flavonoids in plant resistance: a review. Environ Chem Lett. 4(3):147–157. doi:10.1007/s10311-006-0068-8.

Yue R, Lu C, Han X, Guo S, Yan S, Liu L, Fu X, Chen N, Guo X, Chi H, et al. 2018. Comparative proteomic analysis of maize (Zea mays L.) seedlings under rice black-streaked dwarf virus infection. BMC Plant Biol. 18(1):191. doi:10.1186/s12870-018-1419-x.

Zamioudis C, Pieterse CMJ. 2012. Modulation of Host Immunity by Beneficial Microbes. Mol Plant-Microbe Interactions®. 25(2):139–150. doi:10.1094/MPMI-06-11-0179.

Zhang Z, He H, Yan M, Zhao C, Lei C, Li J, Yan F. 2022. Widely targeted analysis of metabolomic changes of Cucumis sativus induced by cucurbit chlorotic yellows virus. BMC Plant Biol. 22(1):158. doi:10.1186/s12870-022-03555-3.

