## Supplementary material for "Beneficial rhizobacteria and virus infection modulate the soybean metabolome and influence the feeding preferences of the virus vector *Epilachna varivestis*": Pulido_et_al_Beetles_Metabolites_supplementary_20240902.rtf

Supplementary Information includes:
•	Supplementary Methods S1.1-S1.4
•	Supplementary Tables S2.1-S2.4
•	Supplementary Figures S3.1-S3.9
•	Supplementary Video S4
•	Supplementary Files S5.1-S5.6
•	Supplementary References S6
•	

S1. SUPPLEMENTARY METHODS
S1.1. Rhizobacteria and BPMV culture conditions
To prepare inoculum sources for all experiments, cultures of B. japonicum and D. acidovorans were grown in yeast-mannitol broth and stored in 30% glycerol at -80°C. For broth culture inoculations, 50 µl of the bacterial glycerol stock was used to inoculate flasks containing broth media. The flasks were shaken at 25°C for 40 hours (D. acidovorans) or 80 hours (B. japonicum). Before seedling inoculation, the liquid inoculum was adjusted to a cell density of 1×10⁹ cfu/ml. Three-day-old seedlings were inoculated with 1 ml of the bacterial suspension applied to the soil using a sterile pipette. Control plants received the same volume of rhizobacteria-free broth.

BPMV, vectored semi-persistently by Epilachna varivestis (Coccinellidae) and Ceratoma trifurcata (Chrysomelidae), causes symptoms like stunted growth, leaf mottling and distortion, and mottling on pods (Giesler et al. 2002). The BPMV strain used was collected from Ohio soybeans, purified to isolate BPMV virions, and then mechanically re-inoculated to generate infected tissue (courtesy of Dr. Peg Redinbaugh). To infect soybean plants, one-week-old seedlings at the V1 stage (post-bacterial inoculation) were dusted with carborundum powder and mechanically inoculated by rubbing leaves with a buffer solution (0.1 M potassium phosphate buffer) containing ground virus-infected tissue. Mock-inoculated plants received the same mechanical treatment with virus-free buffer. Plants were visually monitored for symptom development, and infection was confirmed by ELISA at the conclusion of each experiment.

S1.2. Plant growth and experimental design
Starting from the V1 stage until harvest, plants received 50 mL of a diluted, modified Hoagland's nutrient solution (Dean et al. 2014; Pulido et al. 2019) three times per week. Control and Da plants received a nutrient solution containing the following nitrogen concentrations (in µM): 5001.008 NO₃⁻ and 1675.977 NH₄⁺. For Bj and Bj+Da plants, NO₃⁻ and NH₄⁺ were replaced with K₂SO₄ at equivalent concentrations. Despite these differences in fertilizer composition, soybean plants associated with B. japonicum have been shown to maintain similar foliar nitrogen levels across various nitrogen treatments, enabling a direct comparison between rhizobia-inoculated and control plants (Dean et al. 2014).
In a previous study, we utilized this nitrogen fertilization regime in combination with inoculation by commercially available rhizobial strains, using the same soybean cultivar (Williams 82) and the same environmental chamber setup (Dean et al. 2014). The results indicated that soybean plants maintained similar foliar nitrogen levels, and shoot biomass data suggested that rhizobia inoculation compensated for reduced nitrogen input. Additionally, the presence of a second bacterial colonizer, D. acidovorans, further enhanced plant growth. This approach to supplementing nutrient solution for non-rhizobium inoculated plants has also been successfully employed in similar studies (Juge et al. 2012; Silva et al. 2013; Dean et al. 2014; Kontopoulou et al. 2015).
All plants were grown in an insect-free chamber maintained at 25°C (day) and 23°C (night) under a 16:8 light/dark cycle with 70% relative humidity. At approximately one week of age, half of the plants from each rhizobacteria treatment were manually rub-inoculated with BPMV using a 0.1 M potassium phosphate buffer solution and carborundum powder. The remaining plants received a mock inoculation with virus-free buffer. Virus presence was confirmed using a commercial immunological test (Agdia).
S1.3. Metabolite extraction and analysis
Leaf samples were extracted with 1.0 mL of 80% methanol containing 18 µl/mL umbelliferone as the internal standard for LC-MS analysis. The mixture was vortexed and incubated for 2 hours. After incubation, the samples were centrifuged at 4°C, and 0.5 mL of the supernatant was transferred to a new vial and stored at -20°C for subsequent LC-MS analysis. To the remaining mixture, 1.5 mL of chloroform containing 10 µg/mL docosanol (internal standard for the non-aqueous phase) was added. The samples were sonicated and vortexed between two incubation steps at 50°C. Afterward, the samples were allowed to sit at room temperature, followed by the addition of 1.5 mL of HPLC-grade water containing 25 µg/mL ribitol (internal standard for the aqueous phase). The samples were then vortexed and incubated at 50°C for 45 minutes.
Phase separation was achieved by centrifuging at 2900xg for 30 minutes at 4°C. One mL from each phase was collected into separate 2.0 mL autosampler vials. The aqueous phase was dried in a speed vacuum, while the non-aqueous phase was dried under nitrogen.
For further analysis, the non-aqueous phase was resuspended in chloroform and hydrolyzed with 1.25 M HCl in methanol, followed by a 4-hour incubation at 50°C. After incubation, the samples were dried under nitrogen, resuspended in 70 µL of pyridine, and derivatized with 30 µL of MSTFA + 1% TMCS (Sigma-Aldrich). Following 1-hour incubation at 50°C, the samples were transferred to glass inserts for GC-MS analysis. Dried aqueous extracts were resuspended in pyridine containing 15 mg/mL methoxyamine-HCl, vortexed, and sonicated between two incubation steps at 50°C. Aqueous metabolites were then derivatized with 50 µL of MSTFA + 1% TMCS for 1 hour at 50°C before transfer to glass inserts.
Both aqueous and non-aqueous metabolites were analyzed using an Agilent 7890 GC coupled to a 5975 MSD. Compounds were injected at 230°C and separated on an HP-5MS capillary column (30 m × 0.25 mm ID × 0.25 ìm film thickness; Agilent) under the following temperature program: initial hold at 70°C for 5 minutes, followed by a ramp of 5°C/min to a final temperature of 315°C, held for 12 minutes. One microliter of each derivatized aqueous sample was injected using a 2:1 split ratio, while 0.3 µL of each derivatized non-aqueous sample was injected in splitless mode. Helium was used as the carrier gas at a constant flow of 1.0 mL/min. The MS was operated in electron impact mode (70 eV) with the following settings: Transfer line: 250°C (polar phase), 230°C (non-polar phase); Source: 230°C; Quadrupole: 150°C; Mass scan range: 50–650 amu. Deconvolution algorithms were applied to the total ion chromatograms (TICs) using MassHunter Workstation software (B.06.00; Agilent Technologies), and compounds were identified by comparison with the NIST14 spectral library. Quantification was performed relative to internal standards (25 µg/mL ribitol for the aqueous phase and 10 µg/mL docosanol for the non-aqueous phase).
For secondary metabolite analysis, 5 µL of methanol extract was separated by reverse-phase HPLC using a Prominence 20 UFLCXR system (Shimadzu, Columbia, MD) with a Waters BEH C18 column (100 mm x 2.1 mm, 1.7 µm particle size) maintained at 55°C. Separation was achieved with a 20-minute aqueous acetonitrile gradient at a flow rate of 250 µL/min. Solvent A was HPLC-grade water with 0.1% formic acid, and Solvent B was HPLC-grade acetonitrile with 0.1% formic acid. The initial condition of 97% A and 3% B was increased to 45% B at 10 minutes, 75% B at 12 minutes, held at 75% B until 17.5 minutes, and then returned to initial conditions.
The eluate was analyzed on a 5600 TripleTOF using a Duospray™ ion source (AB Sciex, Framingham, MA). The capillary voltage was set at 5.5 kV in positive ion mode and 4.5 kV in negative ion mode, with a declustering potential of 80 V. The mass spectrometer operated in Information Dependent Acquisition (IDA) mode with a 100 ms survey scan from 100 to 1200 m/z, and up to 20 MS/MS product ion scans (100 ms each) per duty cycle with a collision energy of 50 V and a 20 V spread. LC-MS raw files were processed using XCMS online for retention time alignment, automatic integration, and feature detection. Peak intensities were exported for multivariate analysis, and the METLIN database was used to identify key LC-MS features based on MS signatures and tandem mass spectrometry (MS/MS) spectra.
Data integration was performed after normalizing individual databases from each phase, including secondary metabolites (LC-MS phase) and phytohormones, by their respective phase-specific internal standards and initial sample weights. Compound names were coded according to the IDs listed in Supplementary Table S2.3.
S1.4 Transcriptomics and metabolomics data integration
We used Discriminant Analysis of Principal Components (DAPC) (Jombart et al. 2010) to explore data structure and assess group separation in both transcriptomics and metabolomics datasets, particularly between the rhizobacteria and virus treatments.
To identify the important genes and metabolites that change according to the treatment, we used various feature selection techniques. For metabolites, we employed a recursive feature elimination approach (RFE) and multiple linear models (limma), while for the gene matrix, we used RFE, differential expression analysis (DESeq2), and Weighted correlation network analysis (WGCNA). The end result of these analyses is a list of signature genes and metabolites that were combined to obtain an Integrated Transcriptome-Metabolome Signature (ITMS).
We implemented RFE in the caret R package (Kuhn 2008) using a 5-fold cross-validation resampling method and three different machine learning algorithms: random forest (rf, ntrees = 5000), bagged adaptive boosting (adabag, ntrees = 5000), and support vector machines (svm). This technique helped us identify the most important features in the gene and metabolite sets by recursively removing features from the dataset and retraining the model on the remaining features until we obtained a model with the highest possible accuracy. At each step, we evaluated the importance of the remaining features and eliminated the least important feature.
Next, we used the limma R package (Ritchie et al. 2015) to identify significant differences (p-value < 0.05) in metabolite abundance between two given treatments. We fitted a linear model to the metabolite abundance data and used empirical Bayes statistics to calculate the significance of the differences in metabolite abundance between the groups.
The differential gene expression analysis between two given treatments was achieved with DESeq2 (Love et al. 2014) using a threshold of an adjusted p-value < 0.01. DESeq2 identified the genes that were differentially expressed between two or more experimental conditions by fitting a negative binomial model to the read count data and using empirical Bayes shrinkage to estimate the variance of the gene expression changes.
In addition, we used weighted gene co-expression network analysis (WGCNA) (Langfelder and Horvath 2008) to identify co-expressed gene modules and important genes by detecting significant differences of genes across treatment groups within each module. We used a soft threshold (power = 12) based on the approximate scale-free topology and constructed an unsigned gene network to identify modules of strongly correlated genes using Pearson correlation. We did not pre-filter the gene expression data by differential expression, which allowed us to reveal gene sets that may be highly correlated with specific treatments but do not pass the differential expression threshold (Sánchez-Baizán et al. 2022).
Following the feature selection techniques described above (RFE, DESeq2, limma, and WGCNA), we combined the list of signature genes and signature metabolites into an Integrated Transcriptome-Metabolome Signature (ITMS) for downstream network and integrative analysis. To identify metabolic processes affected by the treatments, we performed gene/metabolite set enrichment analysis using the gage Bioconductor package (Luo et al. 2009) and selected significantly up- and downregulated pathways to map the genes and metabolites in the Metabolism and Environmental Information Processing sections of the KEGG PATHWAY database (http://www.genome.jp/kegg/pathway.html). We generated pathway maps showing differentially expressed genes and metabolites for each Metabolism pathway using the “pathview” function (Luo and Brouwer 2013).
To simplify the comparison of treatments and account for the similarities in gene expression and metabolite abundance patterns observed among the rhizobacteria treatments, we created two new categories: one for all uninfected rhizobacteria treatments (Bj, Bj+Da, and Da) and another for all BPMV-infected rhizobacteria treatments. These categories are referred to as "all Bacteria-uninfected" and "all Bacteria-infected", respectively. This allowed us to compare each combined group against the control-uninfected treatment and identify significant differences between the two groups in the various analyses performed.


S2. SUPPLEMENTARY TABLES:
Table S2.1. Factorial design used to evaluate the main effect of rhizobacterial inoculation and BPMV-infection in soybean plants.
	 	Rhizobacteria treatments	
		Control	B. japonicum	D. acidovorans	B. japonicum + D. acidovorans	
BPMV treatments	infected	Control-BPMV	Bj-BPMV	Da-BPMV	Bj+Da-BPMV	
	uninfected	Control-uninfected	Bj-uninfected	Da-uninfected	Bj+Da-uninfected	

Table S2.2. Comparisons tested in the behavioral assays, and statistical analysis for signature gene and compound selection. DEG = differentially expressed genes by deseq2; DAM = differentially accumulated metabolites by limma. Bj = Bradyrhizobium japonicum, Da = Delftia acidovorans. 

id	Virus treatment	Choice 1	Choice 2	Adult beetles dual-choice test	DEG	DAM	WGCNA	
a.uninfected	Uninfected 	Bj	Control	x	x	x	 	
b.uninfected	Uninfected	Bj+Da	Control	x	x	x	 	
c.uninfected	Uninfected	Da	Control	x	x	x	 	
d.uninfected	Uninfected	Bj+Da	Bj	x	x	x	 	
a.BPMV	Infected	Bj	Control	x	x	x	 	
b.BPMV	Infected	Bj+Da	Control	x	x	x	 	
c.BPMV	Infected	Da	Control	x	x	x	 	
d.BPMV	Infected	Bj+Da	Bj	x	x	x	 	
a.mixed	Mixed	Control infected	control uninfected	 	x	x	 	
b.mixed	Mixed	Bj+Da infected	Bj+Da uninfected 	 	x	x	 	
c.mixed	Mixed	Bj infected	Bj uninfected	 	x	x	 	
d.mixed	Mixed	Bj infected	control uninfected	 	x	x	 	
e.mixed	Mixed	Bj+Da infected	control uninfected	x	x	x	 	
f.mixed	Mixed	Da infected	control uninfected	 	x	x	 	
g.mixed	Mixed	(Bj, Bj+Da, Da) uninfected	control uninfected	 	x	x	x	
h.mixed	Mixed	(Bj, Bj+Da, Da) infected	control uninfected	 	x	x	x	
i.mixed	Mixed	(Bj, Bj+Da, Da) infected	(Bj, Bj+Da, Da) BPMV uninfected	 	x	x	x	


Table S2.3. Compound IDs with KEGG and HMDB codes when available.
ID	Compound Name	KEGG	HMDB	Class	
N3	glycolic acid	C00160	HMDB00115	carboxylic acid	
N6	benzoic Acid	C00180	HMDB01870	carboxylic acid	
N11	glycerol	C00116	HMDB00131	sugar alcohol	
N13	maltol	C11918	HMDB30776	pyranone	
N22	beta-L-Galactopyranoside	 	 	carbohydrate	
N23	L-rhamnose	C00507	HMDB00849	monosaccharide	
N26	methyl xylopyranoside	 	 	carbohydrate	
N29	threonolactone	 	HMDB00940	carboxylic acid	
N31	phosphoric acid	C00009	HMDB02142	inorganic acid	
N33	phosphoric acid	C00009	HMDB02142	inorganic acid	
N35	phosphoric acid	C00009	HMDB02142	inorganic acid	
N37	beta-D-Galactofuranoside	 	 	carbohydrate	
N39	methyl galactoside	 	HMDB29965	carbohydrate	
N42	3,7,11,15-Tetramethyl-2-hexadecen-1-ol	 	 	alcohol	
N43	citronellyl valerate	 	HMDB37229	fatty acid	
N45	D-Mannose	C00159	HMDB00169	monosaccharide	
N47	hexadecanoic acid, methyl ester	C16995	HMDB61859	fatty acid	
N49	iduronic acid	C06472	HMDB02704	carbohydrate	
N52	11,14-Octadecadienoic acid, methyl ester	C01595	HMDB00673	fatty acid	
N57	methyl stearate	 	HMDB34154	fatty acid	
N59	myo-Inositol	C00137	HMDB00211	carbohydrate	
N61	methyl 2-hydroxyhexadecanoate	 	 	fatty acid	
N65	eicosanoic acid	C06425	HMDB02212	fatty acid	
N68	glyceryl-glycoside	 	 	carbohydrate	
N70	D-Galactopyranoside	C03619	 	glucosinolate	
N75	docosanoic acid	C08281	HMDB00944	fatty acid	
N79	methyl 2-hydroxydocosanoate	 	 	fatty acid	
N80	tetracosanoic acid, methyl ester	C08320	HMDB02003	fatty acid	
N81	maltose	C00208	HMDB00163	carbohydrate	
N90	1-hexacosanol	C08381	 	fatty alcohol	
N95	alpha-D-lactose	C00243	HMDB00186	carbohydrate	
N103	beta-sitosterol	C01753	HMDB00852	lipid	
N104	beta-amyrin	C08616	HMDB36658	terpenoid	
P2	propanoic acid	C00163	HMDB00237	carboxylic acid	
P3	pinacol	 	 	alcohol	
P7	2-pentanol	C16834	HMDB31599	alcohol	
P8	acetoin	C00466	HMDB03243	Acyloins	
P9	pentenoic acid	C00803	HMDB00892	carboxylic acid	
P12	3-furoic acid	C01546	HMDB00444	carboxylic acid	
P13	oxalic acid	C00209	HMDB02329	carboxylic acid	
P18	1-octanol	C00756	HMDB01183	fatty alcohol	
P19	propanedioic acid	C00383	HMDB00691	carboxylic acid	
P22	urea	C00086	HMDB00294	organic acid	
P26	1-phenyl-1,2-ethanediol	 	 	alcohol	
P27	dopamine	C03758	HMDB00073	amine	
P32	niacin	C00253	HMDB01488	carboxylic acid	
P34	2-butenoic acid	C01771	HMDB10720	fatty acid	
P35	glycine	C00037	HMDB00123	amino acid	
P36	butanedioic acid	C00042	HMDB00254	carboxylic acid	
P38	glyceric acid	C00258	HMDB00139	carbohydrate	
P39	2-butenedioic acid	C01384	HMDB00176	carboxylic acid	
P41	serine	C00065	HMDB00187	carboxylic acid	
P42	erythrono-1,4-lactone	 	HMDB00349	carbohydrate	
P46	L-threonine	C00188	HMDB00167	amino acid	
P47	hexanoic acid	C01585	HMDB00535	fatty acid	
P49	beta-alanine	C00099	HMDB00056	amino acid	
P51	aspartic acid	C00049	HMDB00191	amino acid	
P52	pyruvic acid	C00022	HMDB00243	carboxylic acid	
P53	malic acid	C00711	HMDB00744	carboxylic acid	
P55	arabino-hexos-2-ulose	 	HMDB29932	aldehyde	
P57	L-5-oxoproline	C01879	HMDB00267	amino acid	
P58	4-aminobutanoic acid	C00334	HMDB00112	amino acid	
P60	propanetriol, 2-methyl-	 	 	alcohol	
P61	L-threonic acid	C01620	HMDB00943	amino acid	
P63	pentanedioic acid	C00489	HMDB00661	carboxylic acid	
P64	xylose	C00181	HMDB00098	monosaccharide	
P66	3-hydroxybenzoic acid	C00587	HMDB02466	carboxylic acid	
P73	hexanedioic acid	C06104	HMDB00448	carboxylic acid	
P75	d-ribose	C00121	HMDB00283	monosaccharide	
P78	putrescine	C00134	HMDB01414	amine	
P90	ribonic acid	C01685	HMDB00867	carbohydrate	
P95	shikimic acid	C00493	HMDB03070	alcohol	
P98	cadaverine	C01672	HMDB02322	amine	
P100	citric acid	C00158	HMDB00094	carboxylic acid	
P106	indolin-2-one	C12312	HMDB61918	indol	
P110	D-pinitol	C03844	HMDB34219	alcohol	
P113	D-fructose	C02336	HMDB00660	monosaccharide	
P118	D-fructose	C02336	HMDB00660	monosaccharide	
P120	levoglucosan	 	HMDB00640	monosaccharide	
P123	D-talose	C06467	 	carbohydrate	
P125	4-coumaric acid	C00811	HMDB02035	carboxylic acid	
P127	D-allose	C01487	HMDB01151	monosaccharide	
P129	D-mannitol	C00392	HMDB00765	carbohydrate	
P130	D-glucitol	C00794	HMDB00247	carbohydrate	
P138	D-gluconic acid	C00257	HMDB00625	carbohydrate	
P140	palmitic acid	C00249	HMDB00220	fatty acid	
P144	D-glucopyranoside	C00738	HMDB62170	monosaccharide	
P145	galactaric acid	C00879	HMDB00639	carbohydrate	
P148	ferulic acid	C01494	HMDB00954	carboxylic acid	
P151	caffeic acid	C01481	HMDB01964	carboxylic acid	
P154	sedoheptulose	C02076	HMDB03219	monosaccharide	
P155	galactose	C00984	HMDB00143	monosaccharide	
P171	D-glucuronic acid	C00191	HMDB00127	carboxylic acid	
P172	2-hydroxymandelic acid, ethyl ester, di-TMS	C11527	HMDB00822	carboxylic acid	
P180	mannobiose	C20861	HMDB29933	carbohydrate	
P192	sucrose	C00089	HMDB00258	carbohydrate	
P198	D-xylopyranose	C00181	HMDB00098	monosaccharide	
P203	D-trehalose	C01083	HMDB00975	carbohydrate	
P207	D-myo-Inisitol	C00137	HMDB00211	alcohol	
P227	dihydroxymalonic acid	C00830	HMDB31522	carboxylic acid	
P235	cellobiose	C06422	HMDB00055	carbohydrate	
P237	D-glucose	C00031	HMDB00122	monosaccharide	
S2	isoquercitrin	C05623	HMDB37362	flavonoid	
S3	L-beta-aspartyl-L-phenylalanine	 	HMDB11167	amino acid	
S5	eriocitrin	C09732	HMDB05811	flavonoid	
S6	kaempferol	C05903	HMDB05801	flavonoid	
S7	formononetin 7-O-rutinoside	C00858	HMDB05808	flavonoid	
S8	pelargonidin 3-gentiotrioside	C05904	HMDB03263	flavonoid	
S9	rubrofusarin	C09047	HMDB34569	glycoside	
S10	rutin	C05625	HMDB03249	flavonoid	
S12	cyanidin 3-rhamnoside 5-glucoside	 	HMDB37993	flavonoid	
S13	cyanidin 3-glucogalactoside	 	HMDB31468	flavonoid	
S14	pelargonidin 3-galactoside-5-glucoside	 	HMDB41170	flavonoid	
S15	biotin-XX hydrazide	C00120	HMDB00030	amine	
S17	caffeic acid 3-O-glucuronide	C01481	HMDB41705	carboxylic acid	
S18	pelargonidin 3-sophoroside 5-glucoside	 	HMDB33687	flavonoid	
S20	luteolin	C01514	HMDB05800	flavonoid	
S22	soyasaponin II	C12081	HMDB34650	lipid	
S23	pratensin A	 	 	flavonoid	
S29	alpha-calendic acid	 	HMDB30962	lipid	
S32	peonidin 3-lathyroside	 	HMDB41171	flavonoid	
S33	ribose-1-arsenate	 	HMDB12285	monosaccharide	
S35	carnosifloside III	C08791	 	terpenoid	
S36	unkown33	 	 	peptide	
S38	1,6-digalloyl-beta-D-glucopyranose	 	HMDB39179	flavonoid	
S39	petunidin 3-rhamnoside 5-glucoside	 	HMDB38099	flavonoid	
S40	chlortetracycline	C06571	HMDB14401	tetracycline	
S43	kuwanon L	 	HMDB30121	flavonoid	
S45	peonidin 3-galactoside-5-glucoside	 	HMDB29210	flavonoid	
S46	petunidin 3-galactoside	 	HMDB38093	flavonoid	
S50	soyasaponin A3	 	HMDB38608	flavonoid	
S52	spinacetin 3-gentiobioside	 	HMDB37469	flavonoid	
S53	Lys Lys Thr Trp	 	 	peptide	
S59	soyasaponin I	C08983	HMDB34649	flavonoid	
S65	PG(16:1(9Z)/0:0)	 	HMDB10571	lipid	
S73	irilone 4'-O-glucoside	 	HMDB33818	flavonoid	
S78	2-O-caffeoylhydroxycitric acid	 	HMDB40572	carboxylic acid	
S82	soyasaponin bg	 	HMDB38721	flavonoid	
S87	2-O-caffeoylglucarate	C03062	HMDB40572	carboxylic acid	
S89	sulindac sulfone	 	HMDB60620	carboxylic acid	
S90	D-ribulose	C00309	HMDB00621	monosaccharide	
S91	Lys Leu Glu Ala Thr	 	 	peptide	
S94	peonidin 3-rutinoside-5-glucoside	 	 	flavonoid	
S95	malvidin 3-sophoroside 5-glucoside	 	HMDB38032	flavonoid	
S97	phosphatidylinositol	C01194	HMDB09807	lipid	
S99	Met Arg Arg Val	 	 	peptide	
S100	araliasaponin III	 	HMDB36477	glycoside	
S120	indomethacin N-octyl amide	 	 	amide	
cisOPDA	12-oxo-phytodienoic acid	C01226	 	phytohormone	
ABA	abscisic acid	C15970	HMDB36093	phytohormone	
S86	gamma-linolenic acid	C06426	HMDB03073	phytohormone	
cisJA	jasmonic acid	C08491	HMDB32797	phytohormone	
LA	linoleic acid	C01595	HMDB00673	phytohormone	
N54	linolenic acid	C06427	HMDB01388	phytohormone	
OA	oleic acid	C00712	HMDB00207	phytohormone	
SA	salicylic acid	C00805	HMDB01895	phytohormone	
S19	tuberonic acid glucoside	C08558	 	phytohormone	
N5	compound_4	 	 	unknown	
N16	compound_7	 	 	unknown	
N24	compound_24	 	 	unknown	
N25	compound_27	 	 	unknown	
N27	compound_32	 	 	unknown	
N32	Unknown N32	 	 	unknown	
N40	Unknown N40	 	 	unknown	
N41	compound_41	 	 	unknown	
N44	compound_43	 	 	unknown	
N50	compound_61	 	 	unknown	
N53	compound_58	 	 	unknown	
N56	compound_46	 	 	unknown	
N58	compound_48	 	 	unknown	
N62	compound_74	 	 	unknown	
N63	compound_76	 	 	unknown	
N64	compound_77	 	 	unknown	
N66	compound_83	 	 	unknown	
N67	compound_85	 	 	unknown	
N69	compound_81	 	 	unknown	
N71	compound_87	 	 	unknown	
N72	compound_53	 	 	unknown	
N74	compound_89	 	 	unknown	
N76	compound_93	 	 	unknown	
N77	compound_94	 	 	unknown	
N78	compound_95	 	 	unknown	
N83	compound_100	 	 	unknown	
N84	compound_102	 	 	unknown	
N85	Unknown N85	 	 	unknown	
N86	compound_106	 	 	unknown	
N87	compound_91	 	 	unknown	
N89	compound_109	 	 	unknown	
N93	compound_103	 	 	unknown	
N96	compound_118	 	 	unknown	
N98	compound_119	 	 	unknown	
N99	compound_108	 	 	unknown	
N105	compound_125	 	 	unknown	
P1	compound_1	 	 	unknown	
P10	compound_6	 	 	unknown	
P14	compound_7	 	 	unknown	
P20	compound_16	 	 	unknown	
P23	compound_18	 	 	unknown	
P37	compound_32	 	 	unknown	
P40	compound_35	 	 	unknown	
P44	compound_30	 	 	unknown	
P56	compound_39	 	 	unknown	
P59	compound_42	 	 	unknown	
P65	compound_49	 	 	unknown	
P67	compound_46	 	 	unknown	
P69	compound_47	 	 	unknown	
P70	compound_52	 	 	unknown	
P71	compound_53	 	 	unknown	
P76	compound_71	 	 	unknown	
P77	compound_60	 	 	unknown	
P79	compound_57	 	 	unknown	
P88	unknown P88	 	 	unknown	
P93	compound_75	 	 	unknown	
P102	compound_69	 	 	unknown	
P107	etoxeridine	 	 	unknown	
P111	compound_84	 	 	unknown	
P112	compound_85	 	 	unknown	
P131	compound_101	 	 	unknown	
P135	compound_116	 	 	unknown	
P136	compound_108	 	 	unknown	
P139	compound_120	 	 	unknown	
P142	compound_114	 	 	unknown	
P147	compound_119	 	 	unknown	
P149	compound_129	 	 	unknown	
P152	compound_123	 	 	unknown	
P153	compound_103	 	 	unknown	
P156	compound_128	 	 	unknown	
P157	compound_137	 	 	unknown	
P161	compound_141	 	 	unknown	
P162	compound_133	 	 	unknown	
P163	compound_110	 	 	unknown	
P174	compound_144	 	 	unknown	
P175	compound_117	 	 	unknown	
P176	unknown N176	 	 	unknown	
P177	compound_122	 	 	unknown	
P178	compound_150	 	 	unknown	
P181	compound_153	 	 	unknown	
P182	compound_125	 	 	unknown	
P183	compound_126	 	 	unknown	
P184	compound_154	 	 	unknown	
P185	unknown P185	 	 	unknown	
P186	unknown P186	 	 	unknown	
P187	compound_159	 	 	unknown	
P193	compound_165	 	 	unknown	
P194	compound_166	 	 	unknown	
P196	compound_168	 	 	unknown	
P197	compound_135	 	 	unknown	
P199	compound_170	 	 	unknown	
P204	compound_176	 	 	unknown	
P205	compound_178	 	 	unknown	
P206	compound_179	 	 	unknown	
P211	compound_187	 	 	unknown	
P219	compound_152	 	 	unknown	
P226	compound_209	 	 	unknown	
P228	unknown P228	 	 	unknown	
P236	compound_221	 	 	unknown	
P238	unknown P238	 	 	unknown	
P239	compound_225	 	 	unknown	
P240	compound_174	 	 	unknown	
P241	compound_175	 	 	unknown	
P243	compound_227	 	 	unknown	
P244	compound_228	 	 	unknown	
P246	compound_230	 	 	unknown	
S24	unknown S24	 	 	unknown	
S25	unknown 74	 	 	unknown	
S26	unknown 97	 	 	unknown	
S28	unknown 96	 	 	unknown	
S34	unknown 71	 	 	unknown	
S42	unknown 77	 	 	unknown	
S48	unknown 78	 	 	unknown	
S56	unknown S56	 	 	unknown	
S57	unknown 84	 	 	unknown	
S62	unknown 76	 	 	unknown	
S63	unknown 93	 	 	unknown	
S69	unknown 80	 	 	unknown	
S72	unknown 46	 	 	unknown	
S74	unknown 70	 	 	unknown	
S76	unknown 85	 	 	unknown	
S77	unknown 64	 	 	unknown	
S79	unknown 48	 	 	unknown	
S88	unknown 79	 	 	unknown	
S96	unknown 67	 	 	unknown	
S98	unknown 60	 	 	unknown	
S101	unknown 55	 	 	unknown	
S102	unknown 69	 	 	unknown	
S103	alhagidin	 	 	unknown	
S105	unknown 81	 	 	unknown	
S106	unknown 57	 	 	unknown	
S108	unknown 95	 	 	unknown	
S109	unknown 61	 	 	unknown	
S110	unknown 66	 	 	unknown	
S112	unknown S112	 	 	unknown	
S117	unknown 62	 	 	unknown	
S124	unknown 68	 	 	unknown	
S126	unknown 63	 	 	unknown	
S127	unknown S127	 	 	unknown	
 


Table S2.4. Top six genes for the modules of co-expression (MEs) shown in Fig. 9 
Module	Top six genes	
6	uncharacterized LOC100777483

uncharacterized LOC100805057

chlorophyllide a oxygenase, chloroplastic

geraniol 8-hydroxylase

chlorophyllide a oxygenase, chloroplastic

uncharacterized LOC100812363

	
1	glutamate decarboxylase

uncharacterized LOC100500099

2-methyl-6-phytyl-1,4-hydroquinone methyltransferase, chloroplastic

protein CELLULOSE SYNTHASE INTERACTIVE 1

uncharacterized LOC100798854

nucleobase-ascorbate transporter 6

	
7	serine carboxypeptidase-like

resistance protein MG13

sodium/calcium exchanger NCL2

LEAF RUST 10 DISEASE-RESISTANCE LOCUS RECEPTOR-LIKE PROTEIN KINASE-like 2.1

WRKY DNA -binding domain-containing protein

WRKY transcription factor 17

	


S3. SUPPLEMENTARY FIGURES


Supplementary Figure S3.1. Feeding and foraging preferences of adult beetles. The activity of five adult beetles was recorded inside the tent. Two plants from different treatments were placed inside a fine-mesh cage.


Supplementary Figure S3.2. Rhizobacteria-induced metabolite changes in uninfected soybean plants. The volcano plot displays the log2 fold change on the x-axis and the negative log10 of the p-value on the y-axis. Significant metabolites (p-value < 0.05) are highlighted as green dots, while non-significant metabolites (p-value > 0.05) are shown as black dots. The top 10 most significant metabolites are labeled in the plot, with compound IDs listed in Table S2.3. 


Supplementary Figure S3.3. Rhizobacteria-induced metabolite changes in BPMV-infected soybean plants. The volcano plot displays the log2 fold change on the x-axis and the negative log10 of the p-value on the y-axis. Significant metabolites (p-value < 0.05) are highlighted as green dots, while non-significant metabolites (p-value > 0.05) are shown as black dots. The top 10 most significant metabolites are labeled in the plot, with compound IDs listed in Table S2.3. 


Supplementary Figure S3.4. BPMV-induced metabolite changes. The volcano plots display the log2 fold change on the x-axis and the negative log10 of the p-value on the y-axis. The top panels show BPMV-induced metabolite changes in rhizobacteria-free and rhizobacteria-inoculated plants, while the bottom panels depict metabolite changes upon mixed treatments. Significant metabolites (p-value < 0.05) are highlighted as green dots, and non-significant metabolites (p-value > 0.05) are shown as black dots. The top 10 most significant metabolites are labeled in the plot, with compound IDs listed in Table S2.3.


Supplementary Figure S3.5. Number of Signature Genes per treatment. The horizontal bar graph displays the number of signature genes identified in various pairwise comparisons of rhizobacteria and BPMV treatments. The x-axis represents the number of signature genes, while each row on the y-axis represents a different pairwise comparison. Bars indicate the number of upregulated genes (in yellow) and downregulated genes (in blue). The integrated Gene Signature (IGS) was identified through deseq analysis, recursive feature elimination, and weighted gene co-expression network analysis.


Supplementary Figure S3.6. Rhizobacteria-induced signature gene profiles of uninfected soybean plants. The volcano plots display the log2 fold change on the x-axis and the negative log10 of the p-value on the y-axis. Names of Signature Genes identified through deseq analysis, recursive feature elimination, and WGCNA are shown. Significant genes (log2 fold change > 1.5 and adjusted p-value < 0.01) are highlighted as green dots, while non-significant genes are shown as black dots. A. Bj vs control. B. Bj+Da vs control. C. Da vs control. D. Bj+Da vs Bj.


Supplementary Figure S3.7. Rhizobacteria-induced gene profiles in BPMV-infected soybean plants. The volcano plots display the log2 fold change on the x-axis and the negative log10 of the p-value on the y-axis. Names of Signature Genes identified through deseq analysis, recursive feature elimination, and WGCNA are shown. Significant genes (log2 fold change > 1.5 and adjusted p-value < 0.01) are highlighted as green dots, while non-significant genes are shown as black dots. A. Bj vs control. B. Bj+Da vs control. C. Da vs control. D. Bj+Da vs Bj.


Supplementary Figure S3.8.  Soybean gene profiles upon rhizobacteria inoculation and BPMV infection. The volcano plots display the log2 fold change on the x-axis and the negative log10 of the p-value on the y-axis. Names of Signature Genes identified through deseq analysis, recursive feature elimination, and WGCNA are shown. Significant genes (log2 fold change > 1.5 and adjusted p-value < 0.01) are highlighted as green dots, while non-significant genes are shown as black dots. A. Bj vs control. B. Bj+Da vs control. C. Da vs control. D. Bj+Da vs Bj.


Supplementary Figure S3.9. KEGG pathways identified through the analysis of fold changes in signature genes and metabolites in soybean plants induced by rhizobacteria and BPMV infection. Pathway perturbations were analyzed across three contrasts: A) all uninfected rhizobacteria treatments compared to control uninfected, B) all rhizobacteria and BPMV-infected treatments compared to control uninfected, and C) all rhizobacteria and BPMV-infected treatments compared to all uninfected rhizobacteria treatments. Each graph represents a significantly perturbed pathway, where most of the signature genes and compounds were either upregulated (top) or downregulated (bottom). Note that a KEGG node may represent multiple genes with similar functions. Additional significant pathways can be found in the supplementary file S5.6 deposited in the ETHZ repository [xxxxx]. all Bacteria = B. japonicum, B. japonicum + D. acidovorans, and D. acidovorans. 
S4. SUPPLEMENTARY VIDEO 
Example of a dual-choice assay demonstrating the foraging behavior of adult beetles. Two plants from different treatments were placed inside a fine-mesh cage, and five beetles were released to forage freely for 24 hours. This setup illustrates beetle preference and feeding behavior under varying experimental conditions.
S5. SUPPLEMENTARY FILES
Supplementary file S5.1. “Supplementary_file_S5.1_results_limma_20230413.xlsx”
This Excel file contains ranked compounds resulting from the linear model fit using the R function limma::topTable() for multiple contrasts. The file includes several sheets, with each sheet representing the results of the limma analysis for the specific pair-wise comparisons shown in Figure 7 and Supplementary Figures S3.2–S3.4.
Key information in the file includes:
Compound Identification: The compound ID, classification, and additional identifiers, such as KEGG and Human Metabolome Database (HMDB) IDs, where available.
Statistical Results: The log fold change (logFC) and p-values from the t-test for each comparison.
 
Supplementary file S5.2. “Supplementary_file_S5.2_results_deseq3_20230413.xlsx”
Summary of differential expression analysis results from DESeq2. 
This file contains multiple sheets, with each sheet presenting results for the specific pair-wise comparisons shown in Figure 8 and Supplementary Figures S3.6–S3.8. Each sheet includes a table with the following columns:
•	id: The identifier for each gene or feature in the dataset.
•	Entrez_id: the entrez id for each gene.
•	symbol: the symbol for each gene.
•	genename: the corresponding gene name when available.
•	baseMean: The mean of normalized counts across all samples for each gene or feature.
•	log2FoldChange: The log2 fold change in expression levels between two conditions (treatment1 vs treatment2). Positive values indicate upregulation in treatment1, while negative values indicate downregulation.
•	pvalue: The p-value associated with the Wald statistic, indicating the likelihood of the observed data under the null hypothesis of no differential expression.
•	padj: The adjusted p-value, accounting for multiple testing correction (e.g., Benjamini-Hochberg). This value represents the probability of false positives, with smaller values indicating more significant results.
 
Supplementary file S5.3. “Supplementary_file_S5.3_wgcna_gene_to_module_membership_20230413.xlsx”
Module membership and position of hub genes identified from the weighted gene co-expression network analysis (WGCNA). The table lists gene ID, Entrez ID, gene symbol, gene name, module membership, and position for each hub gene. The position of the gene indicates its relative importance within the module.
 
Supplementary file S5.4. “Supplementary_file_S5.4_results_limma_wgcna_20230413.xlsx”
Differential gene expression analysis was performed within each module identified by Weighted Gene Co-Expression Network Analysis (WGCNA).
The analysis was conducted across three contrasts:
1)	All rhizobacteria uninfected treatments vs. control uninfected.
2)	All rhizobacteria and BPMV-infected treatments vs. control uninfected.
3)	All rhizobacteria and BPMV-infected treatments vs. all rhizobacteria uninfected treatments.
The table includes the following information for each differentially expressed gene:
•	Gene ID
•	Treatment group names
•	Module ID and module color (assigned by WGCNA)
•	Number of genes in the module (n)
•	Log2 fold change (logFC)
•	T-statistic (t)
•	P-value (P.Value)
•	Adjusted p-value (adj.P.Val)
 
Supplementary file S5.5. “Supplementary_file_S5.5_gage_pathways_result_20230413.xlsx”
Significant enriched pathways from gene/metabolite enrichment analysis were identified across multiple contrasts, including comparisons between rhizobacteria for uninfected or BPMV-infected plants and for mixed virus treatments. Some of these comparisons are depicted in Figure 9.
Table Columns:
•	Regulation: Indicates whether the pathway is upregulated or downregulated.
•	gmx_id: The KEGG pathway identifier.
•	Path: Name of the KEGG pathway.
•	Category1 and Category2: Classification categories of the pathway.
•	p.geomean: Geometric mean of the p-values of the genes in the gene set. This provides an overall measure of the pathway's significance, reflecting the combined effect of all genes in the set.
•	stat.mean: Mean log2 fold change of the genes in the gene set, indicating the overall direction and magnitude of gene expression changes (upregulated or downregulated).
•	p.val: The unadjusted p-value for the gene set, calculated using a hypergeometric test, which measures the likelihood of observing the given number of genes in the set by chance.
•	q.val: Adjusted p-value (or False Discovery Rate, FDR) of the gene set, corrected for multiple testing using the Benjamini-Hochberg method.
•	set.size: The number of genes in the gene set.
•	exp1: Enrichment score for each gene/metabolite set, representing the degree of over-representation of the genes/metabolites in the ranked list, relative to what would be expected by chance. Scores range from 0 to 1, with higher scores indicating greater over-representation. A score of 1 means all genes/metabolites in the set appear at the top of the ranked list, while a score of 0 means the set is randomly distributed in the list.

Supplementary file S5.6. “Supplementary_file_S5.6_KEGG_pathways.zip”
This zip file contains KEGG pathway graphs for each of the pair-wise comparisons in the table below. Each graph depicts a significantly perturbed pathway, where the majority of the signature genes and compounds were either upregulated or downregulated. The KEGG pathways maps are based on the list of signature genes and metabolites used to derive an Integrated Transcriptome-Metabolome Signature (ITMS). The folder names are coded based on the specific pairwise comparison between treatment 1 and treatment 2.

folder name                	 virus   	 treatment1       	 treatment2	
res.a.uninfected   	 uninfected 	 Bj           	 control	
res.b.healthy      	 uninfected 	 Bj+Da        	 control	
res.a.BPMV         	 BPMV      	 Bj           	 control	
res.b.BPMV         	 BPMV      	 Bj+Da        	 control	
res.c.BPMV         	 BPMV      	 Da           	 control	
res.d.BPMV         	 BPMV      	 Bj+Da        	 Bj	
res.a.mixed        	 mixed    	 control-BPMV   	 control-uninfected	
res.b.mixed        	 mixed    	 Bj+Da-BPMV    	 Bj+Da-uninfected	
res.c.mixed        	 mixed    	 Bj-BPMV       	 Bj-uninfected	
res.d.mixed        	 mixed    	 Bj-BPMV       	 control-uninfected	
res.e.mixed        	 mixed    	 Bj+Da-BPMV    	 control-uninfected	
res.f.mixed        	 mixed    	 Da-BPMV       	 control-uninfected	
res.g.mixed        	 mixed    	 allBacteria-uninfected 	 control-uninfected	
res.h.mixed        	 mixed    	 allBacteria-BPMV 	 control-uninfected	
res.i.mixed        	 mixed    	 allBacteria-uninfected 	 allBacteria-BPMV	

S6. SUPPLEMENTARY REFERENCES
Dean JM, Mescher MC, Moraes CM. 2014. Plant dependence on rhizobia for nitrogen influences induced plant defenses and herbivore performance. International Journal of Molecular Sciences. 15:1466–1480. doi:10.3390/ijms15011466.
Giesler LJ, Ghabrial SA, Hunt TE, Hill JH. 2002. Bean pod mottle virus: a threat to US soybean production. Plant Disease. 86:1280–1289. doi:10.1094/PDIS.2002.86.12.1280.
Jombart T, Devillard S, Balloux F. 2010. Discriminant analysis of principal components: a new method for the analysis of genetically structured populations. BMC genetics. 11:1–15. doi:10.1186/1471-2156-11-94.
Juge C, Prevost D, Bertrand A, Bipfubusa M, Chalifour FP. 2012. Growth and biochemical responses of soybean to double and triple microbial associations with Bradyrhizobium, Azospirillum and arbuscular mycorrhizae. Applied Soil Ecology. 61:147–157. doi:10.1016/j.apsoil.2012.05.006.
Kontopoulou CK, Giagkou S, Stathi E, Savvas D, Iannetta PPM. 2015. Responses of Hydroponically Grown Common Bean Fed with Nitrogen-free Nutrient Solution to Root Inoculation with N-2-fixing Bacteria. Hortscience. 50:597–602. doi:10.21273/HORTSCI.50.4.597.
Kuhn M. 2008. Building predictive models in R using the caret package. Journal of statistical software. 28:1–26. doi:10.18637/jss.v028.i05.
Langfelder P, Horvath S. 2008. WGCNA: an R package for weighted correlation network analysis. BMC bioinformatics. 9:1–13. doi:10.1186/1471-2105-9-559.
Love MI, Huber W, Anders S. 2014. Moderated estimation of fold change and dispersion for RNA-seq data with DESeq2. Genome biology. 15:1–21. doi:10.1186/s13059-014-0550-8.
Luo W, Brouwer C. 2013. Pathview: an R/Bioconductor package for pathway-based data integration and visualization. Bioinformatics. 29:1830–1831. doi:10.1093/bioinformatics/btt285.
Luo W, Friedman MS, Shedden K, Hankenson KD, Woolf PJ. 2009. GAGE: generally applicable gene set enrichment for pathway analysis. BMC bioinformatics. 10:1–17. doi:10.1186/1471-2105-10-161.
Pulido H, Mauck KE, Moraes CM, Mescher MC. 2019. Combined effects of mutualistic rhizobacteria counteract virus-induced suppression of indirect plant defences in soya bean. Proceedings of the Royal Society B. 286:20190211. doi:10.1098/rspb.2019.0211.
Ritchie ME, Phipson B, Wu D, Hu Y, Law CW, Shi W, Smyth GK. 2015. limma powers differential expression analyses for RNA-sequencing and microarray studies. Nucleic acids research. 43:47–47. doi:10.1093/nar/gkv007.
Sánchez-Baizán N, Ribas L, Piferrer F. 2022. Improved biomarker discovery through a plot twist in transcriptomic data analysis. BMC biology. 20:1–26. doi:10.1186/s12915-022-01398-w.
Silva LR, Pereira MJ, Azevedo J, Mulas R, Velazquez E, González-Andrés F, Valentão P, Andrade PB. 2013. Inoculation with Bradyrhizobium japonicum enhances the organic and fatty acids content of soybean (Glycine max (L.) Merrill) seeds. Food chemistry. 141:3636–3648. doi:10.1016/j.foodchem.2013.06.045.
